# Modulation of Human Frontal Midline Theta by Neurofeedback: A Systematic Review and Quantitative Meta-Analysis

**DOI:** 10.1101/2023.11.10.566628

**Authors:** Maria Pfeiffer, Andrea Kübler, Kirsten Hilger

**Affiliations:** Institute of Psychology, Department of Psychology I, Würzburg University, Marcusstr. 9-11, Würzburg D-97070, Germany

**Keywords:** Neurofeedback, Frontal Midline Theta, EEG, Executive Functions, Meta-Analysis, Review

## Abstract

Human brain activity consists of different frequency bands associated with varying functions. Oscillatory activity of frontal brain regions in the theta range (4-8Hz) is linked to cognitive processing and can be modulated by neurofeedback - a technique where participants receive real-time feedback about their brain activity and learn to modulate it. However, criticism of this technique evolved, and high heterogeneity of study designs complicates a valid evaluation of its effectiveness. This meta-analysis provides the first systematic overview over studies attempting to modulate frontal midline theta with neurofeedback in healthy human participants. Out of 1431 articles screened, 14 studies were eligible for systematic review and 11 for quantitative meta-analyses. Studies were evaluated following the DIAD model and the PRISMA guidelines. A significant across-study effect of medium size (Hedges’ *g* = .66; 95%-CI [−0.62, 1.73]) with substantial between-study heterogeneity (Q(16) = 167.43, *p* < .0001) was observed and subanalysis revealed effective frontal midline theta upregulation. We discuss moderators of effect sizes and provide guidelines for future research in this dynamic field.

## 1. Introduction

### 1.1. Frontal Midline Theta (FMT)

Electroencephalography (EEG) measures non-invasively oscillatory electrical brain activity at the scalp level. These oscillations are based on synchronized firing of cortical neurons (i.e., mostly parallelly arranged pyramidal neurons) and the resulting local field potentials. The assessed rhythmic brain activity is commonly categorized into five frequency bands (alpha, beta, gamma, delta and theta) which correlate with different cognitive functions (Müller-Putz, 2020). An increase in the amplitude of a specific frequency band measured with EEG is assumed to be induced by an increase of synchronized neuronal firing in the pace of this band and ultimately as a consequence of stronger demands on associated cognitive functions. The current study focuses on the theta frequency band ranging from 4 to 8 Hz, more specifically, on frontal midline theta (FMT) - a frontally localized kind of this oscillatory brain activity.

#### 1.1.1. Historical Background

Although the first record of these frontal theta oscillations can be dated back to the early 1950s (Arellano and Schwab, 1950; Brazier and Casby, 1952), the first time they are referred to as “frontal midline theta” was noted in the 1970s (Ishihara and Yoshii, 1972). Advancements in the processing of the FMT signal in the late 20th-century contributed markedly to a deeper understanding of its underlying neural mechanism and research in the following years revealed associations between FMT and specific cognitive processes including working memory (Asada et al., 1999; Gevins et al., 1997; Ishii et al., 1999) and episodic memory encoding and retrieval (Klimesch, 1999; Klimesch et al., 2000).

#### 1.1.2. Localization and Function

Frontal midline theta can be best obtained from mid-frontal electrodes at Fz or anterior to Fz (Mitchell et al., 2008) according to the 10-20 system (Acharya et al., 2016; Jasper, 1958). Its source is suggested to lie in the dorsal anterior cingulate cortex (dACC; Cavanagh and Frank, 2014; Mitchell et al., 2008), an anterior subregion of the midcingulate cortex. A large body of research supports dACC-induced FMT as a signal of cognitive control and executive functioning, especially of conflict experience, conflict monitoring and of top-down control (Cavanagh and Shackman, 2015; Mitchell et al., 2008; Nigbur et al., 2011). In line with this assumption, FMT has also been associated with the occurrence of event-related potentials such as the N2 as well as with feedback-, correct-, or error-related negativities (FRN, CRN and ERN; Cavanagh and Frank, 2014).

Nonetheless, it remains challenging to ascribe FMT to a particular cognitive function (Mitchell et al., 2008). While FMT increases with the intake of anxiolytic drugs and shows a negative correlation with state anxiety (Mitchell et al., 2008), positive correlations have also been reported with state (Cavanagh and Shackman, 2015) and trait anxiety (Osinsky et al., 2017; Schmidt et al., 2018). Further cognitive processes that have been associated with FMT include memory formation and memory retrieval (Hsieh and Ranganath, 2014), along with increases in FMT during meditation (Brandmeyer and Delorme, 2018; Cahn and Polich, 2006) and during subjective flow experience (Katahira et al., 2018). Thus, the one key aspect in the elicitation of FMT can be assumed to be internally focused cognition (Cavanagh and Cohen, 2022).

#### 1.1.3. Clinical Relevance

The involvement of FMT in this broad spectrum of cognitive and emotional processes renders it an attractive target for studying its potential as a biomarker for diverse psychological diseases and for clinical intervention. In fact, aberrant FMT oscillations have been observed in diverse clinical subgroups. For example, increased FMT was noticed in patients with anxiety, while diminished FMT was suggested to characterize patients with externalizing or substance abuse disorders (Gilmore et al., 2010; Kamarajan et al., 2015). Decreased theta habituation to a novel sound served as a main predictor for the successful classification of patients Parkinson’s disease into an ON and OFF medication group (Cavanagh et al., 2018), whereas a lower, FMT-related mismatch negativity reliably predicted the indication of schizophrenia (Light et al., 2015), further supporting its role as a neural indicator of clinical diseases.

Based on these and complementary research findings, FMT was also proposed as an attractive target for induced modulation – not only for therapeutic purpose and cognitive or emotional enhancement, but also for a more specific investigation of this oscillatory brain activity itself. To this aim, various approaches were developed during the last decades to externally modulate FMT with transcranial alternating current stimulation (tACS; Chander et al., 2016; Shtoots et al., 2022), transcranial direct current stimulation (tDCS; Choe et al., 2016; Miller et al., 2015) or low intensity focused ultrasound (LITFUS; Forster et al., 2023; Ziebell et al., 2022). In addition to these external tools, means that enable the intrinsic modulation of FMT have also been established, in particular neurofeedback (see Table 1).

**Table 1.**
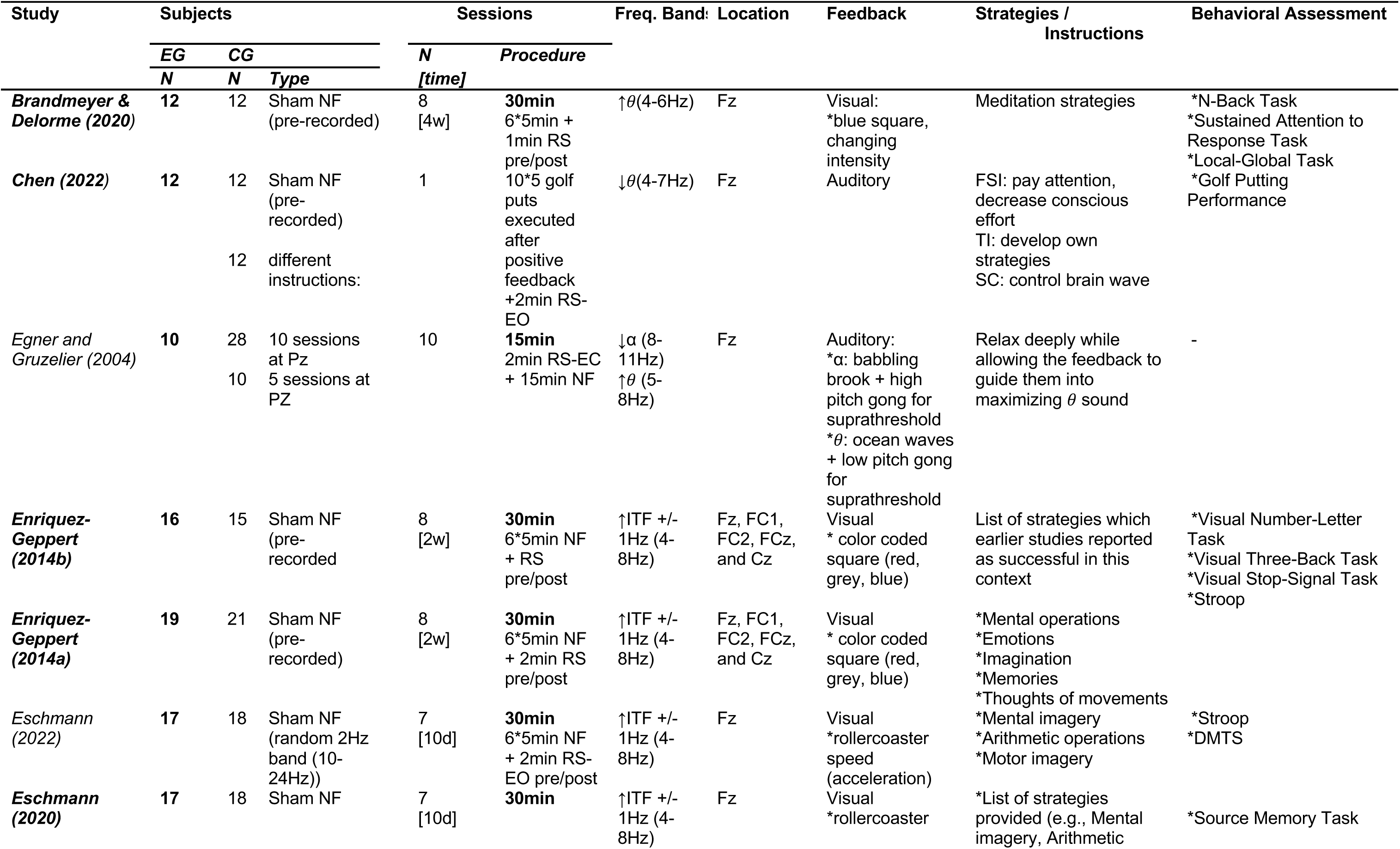
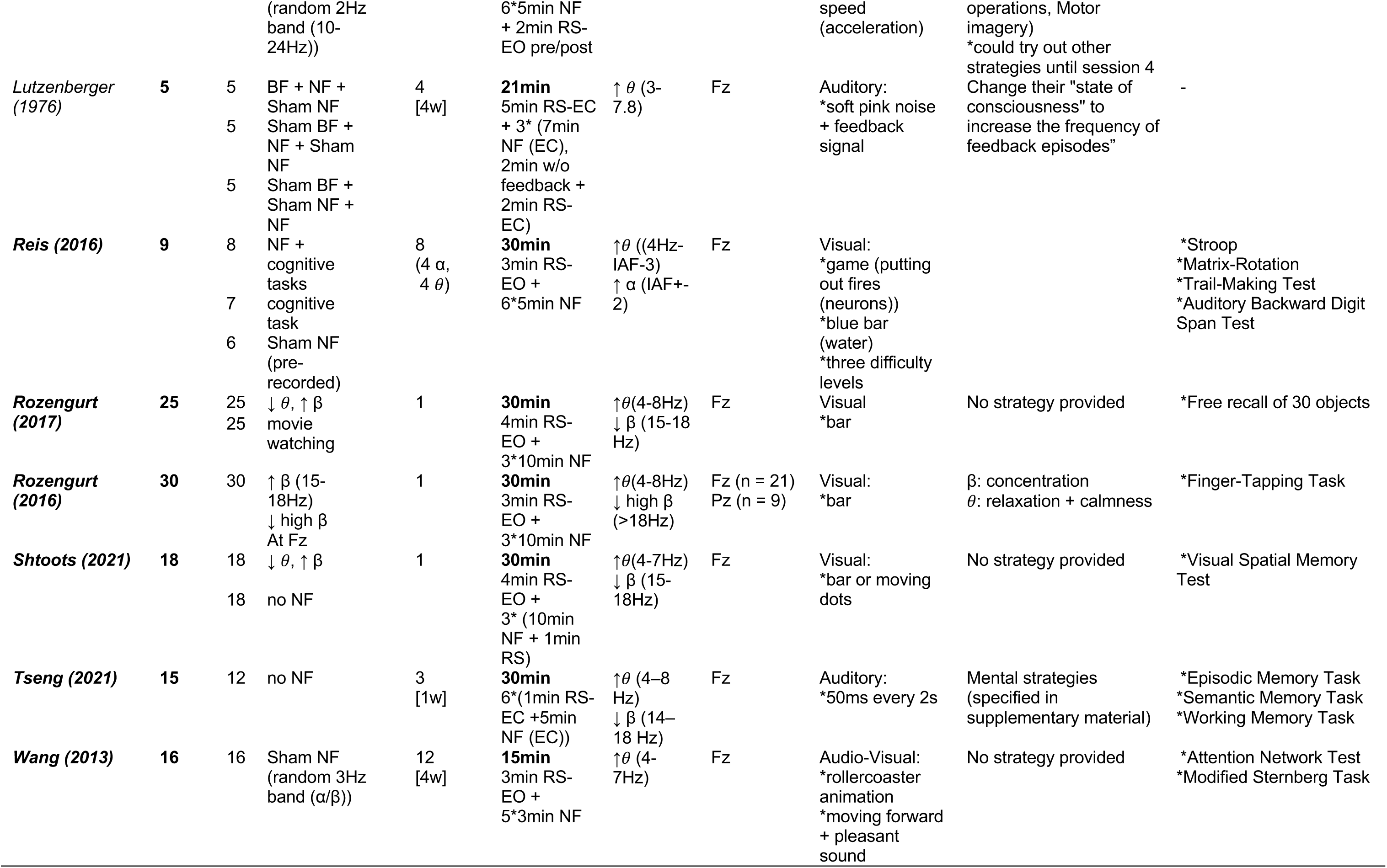
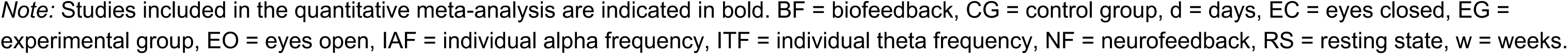
Overview of the studies included in the qualitative review and in the quantitative meta-analysis.

### 1.2. Neurofeedback

Neurofeedback (NF) is biofeedback aiming at the deliberate control of specific brain states. By becoming aware of a specific brain state such as the amplitude of a defined oscillatory brain activity or by being rewarded for it (e.g., by positive feedback), or by both (becoming aware and being rewarded), subjects can learn to deliberately modulate a particular brain activity (Sitaram et al., 2017). Already in 1962, the feasibility of modulating brain activity in humans has been demonstrated (Kamiya, 1962) and more recent investigations suggest that changes in oscillatory activity (Sterman et al., 1970) and plasticity (Ros et al., 2010) induced by neurofeedback can even exceed the period of active training.

#### 1.2.1 Study Design

In therapeutical practice, a typical neurofeedback intervention includes multiple *sessions*. Especially in treatment for ADHD (attention-deficit hyperactivity disorder), protocols with up to 40 sessions are not uncommon (Gevensleben et al., 2012; Meisel et al., 2013; Steiner et al., 2014), while other studies focus on effects of a single neurofeedback session. Each session can be further subdivided into different time segments, so-called *runs*, which allows the participant to rest and to recover. Often, a resting-state EEG is recorded at the beginning and at the end of a session, the former to calibrate the system, the latter to assess the sustainability of the induced effects. While resting state is defined by the absence of an explicit task, the instructions and exact implementation of resting-state measurements differ broadly (Finn, 2021). Some studies also include so-called *transfer runs*, where participants are asked to modulate the signal, but do not receive any feedback. Those runs allow to differentiate between modulations caused by a deliberately internal state change and those merely reflecting the perception of the feedback signal itself. Finally, it is important to note that any neurofeedback approach requires a high temporal resolution for data acquisition and processing to deliver the required real-time feedback. Such feedback can, in principle, be administered via any sensory modality, but auditory and visual stimuli are most commonly used.

#### 1.2.2. Practical Applications

The purpose of neurofeedback interventions is often to improve our basic understanding about the relationships between physiological states and their behavioral correlates, but also to normalize aberrant brain states in psychological disorders (e.g., ADHD; Begemann et al., 2016; Van Doren et al., 2019) or to enhance specific cognitive processes (Viviani and Vallesi, 2021). The resultant neurofeedback-induced changes in neurological symptoms or cognitive capabilities are typically assessed with behavioral tasks and questionnaires.

A common criticism regarding the use of neurofeedback is that studies often exclusively report changes in behavioral outcomes, while descriptions of potential changes in neural measures are lacking, thus, leaving the question unanswered whether the observed behavioral changes are actually driven by the modulated brain states. This doubts the specificity of neurofeedback interventions (Thibault et al., 2017), especially when considering the circumstance that roughly 30% of participants do not express any changes on the neural level (so called non-responders; Alkoby et al., 2018; Allison and Neuper, 2010; Gruzelier, 2014) and that the underlying neurobiological mechanisms of neurofeedback-learning are far from being understood.

#### 1.2.3. Underlying Mechanisms

Previously, neurofeedback-learning was assumed to rely solely on operant conditioning, while today various theories proclaim conscious processing of the feedback signal as a key factor (Sitaram et al., 2017). Also, the exact mechanisms of long-term changes are not yet completely understood. Both, homeostatic (Kluetsch et al., 2014) and Hebbian (Cho et al., 2008; Ros et al., 2013) plasticity after neurofeedback-training were observed (Ros et al., 2014). Most critically, even though neurofeedback has been studied for almost six decades, there is still a lack of definite empirical evidence supporting its efficacy (Thibault et al., 2017). The lack of proper controls, small sample sizes, heterogeneous protocols and contradictory results are frequent and complicate drawing general conclusions. To foster proper study designs and to increase across-study comparability of analysis and results, first guidelines for neurofeedback studies have recently been proposed (Rogala et al., 2016; Ros et al., 2020). Nevertheless, the number of possible design choices remain high and requires a thorough evaluation for any specific neurofeedback application (e.g., such as FMT neurofeedback). Further, the question of what is referred to as “effective” may vary vastly between neurofeedback studies.

### 1.3. FMT Neurofeedback

Neurofeedback has been used to modulate human FMT brain activity (Enriquez-Geppert et al., 2014b; Lutzenberger et al., 1976), with the aim of enhancing cognitive function (Wang and Hsieh, 2013), but also directed at the research of novel therapeutic approaches (Enriquez-Geppert et al., 2016; Lackner et al., 2016; Smit et al., 2023).

However, FMT neurofeedback is accompanied by the same criticism as neurofeedback in general, most importantly, it is not sufficiently clarified whether it is effective at all. Standardized protocols are missing, and the current state of research presents highly heterogeneous study designs, which impede across-study comparison of very heterogeneous findings. What is lacking, so far, is a comprehensive comparison of study parameters and a systematic synthesis of FMT neurofeedback results, to allow for a more empirically based choice of design parameters.

### 1.4. Aim of this Meta-Analysis

We provide a systematic evaluation and meta-analytical investigation of all studies, published until October 2022, attempting to modulate frontal midline theta via neurofeedback in healthy adult participants. To this aim, we systematically compared and evaluated different study design characteristics to derive insights into how these may affect study outcomes. Further, we provide a new nomenclature for distinguishing different types of effectiveness. Overall, we strive for identifying important shortcomings, pressing challenges and the most promising developments with the ultimate goal to provide useful recommendations and guidelines for future research in this dynamic field.

## 2. Methods

The protocol for this study was formally preregistered in the Open Science Framework: https://osf.io/qymhr. Please note that due to initial overestimation of extractable data (fewer studies with more heterogeneous study designs and more missing reports of measurements than expected) it was necessary to deviate in few details from the original protocol, in particular with regard to the quantitative meta-analysis. Any such modifications made to the preregistered protocol are marked.

### 2.1. Study Selection

Studies published until October 2022 were retrieved and selected using the Preferred Reporting Items for Systematic Reviews and Meta-Analyses (PRISMA, Page et al., 2021; see Fig. 1 for a flowchart diagram of the whole study selection procedure). We consulted the electronic databases PubMed, Web of Science, MedRxiv and PsyArXiv and used the following search terms with the following Boolean operators: *“(theta) AND ((neurofeedback) OR (EEG neuromodulation) OR (biofeedback)) NOT (stimulation) NOT (animal) NOT (rodent) NOT (rat)”*. This strategy yielded 1186 results after duplicates were removed. Additionally, Google Scholar and Research Gate were considered with the query *“theta AND (neurofeedback OR EEG neuromodulation OR biofeedback) -stimulation -animal -rodent -rat”,* which led to the inclusion of additional 30 reports. The abstracts and titles of these 1216 papers were screened for the following, primary exclusion criteria: (1) The use of non-human subjects, (2) primary focus on children or adolescents (< 18 years), (3) no inclusion of at least one healthy neurofeedback group and one healthy control group, thus, not allowing to draw direct conclusions about the modulation effect in healthy adults, and (4) no EEG-neurofeedback application or (5) article not available in English or German. This led to the exclusion of 1067 papers.

**Fig. 1.**
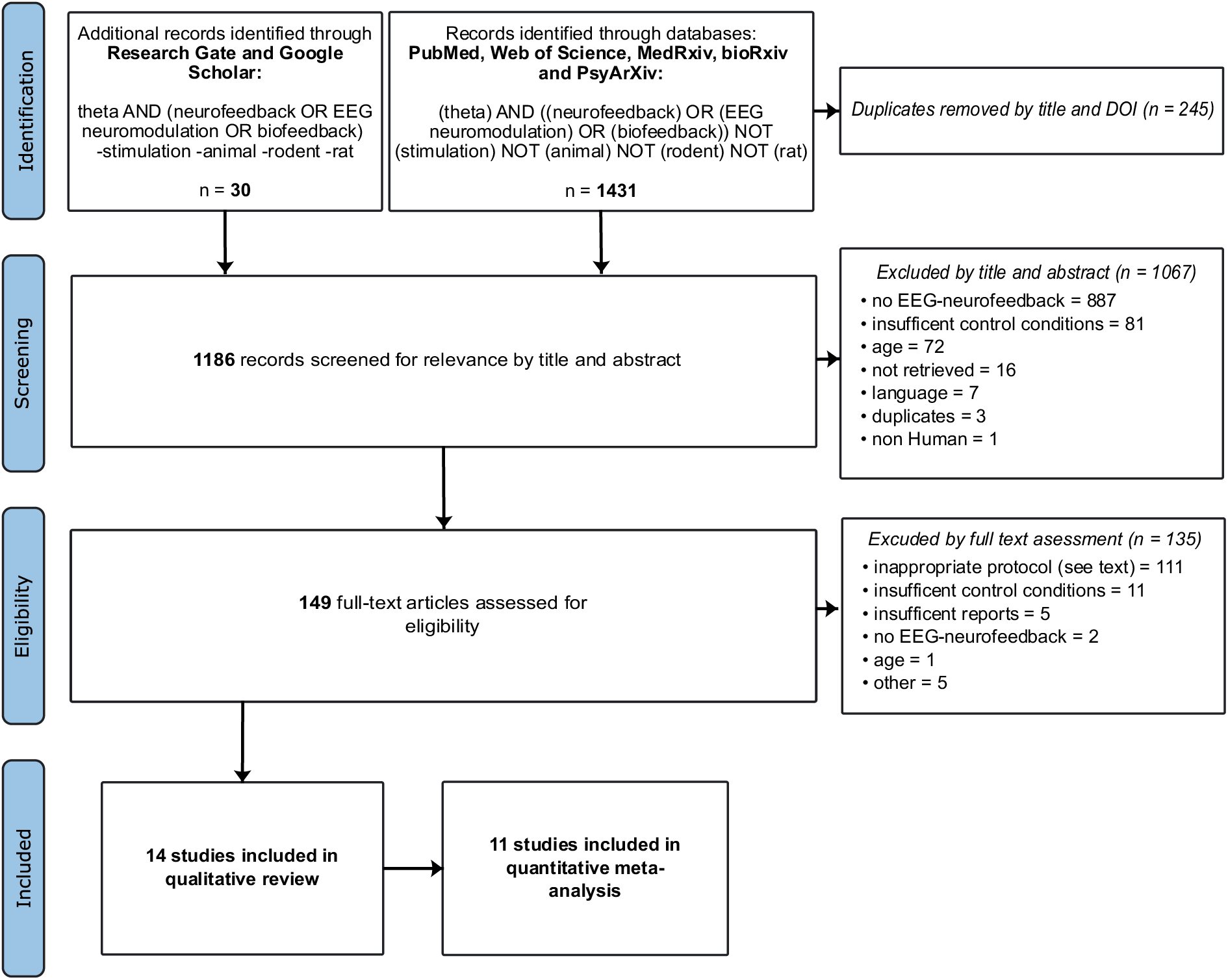
PRISMA flowchart of the study selection process. Exclusion of studies are indicated by italic headers. Of 1431 screened articles, fourteen were included in the qualitative review and eleven in the quantitative meta-analysis.

In a second step, full-texts of the remaining 149 reports were screened for the secondary exclusion criteria: (1) Insufficient reports (e.g., modulation success not reported - only behavioral changes, etc.), (2) no predefined direction of modulation (e.g., modulation protocol based on deviation from a baseline, normative database, or another participant), (3) FMT modulation not intended (feedback not calculated from the frequency of interest, i.e., between 3-9 Hz), or (4) focus on different brain areas as typically associated with FMT, i.e., from frontal midline electrodes (AFz, Fz or FCz) or the dACC. This resulted in a final sample of 14 studies eligible for inclusion in the qualitative review.

### 2.2. Risk of Bias Assessment

To systematically and objectively assess study quality and potential biases in study design, all studies included in the review were rated according to the Study Implementation Assessment Device (Study DIAD; Valentine and Cooper, 2008), which is a general system for evaluating study design quality, can be applied across various fields of research and has been successfully used in other non-clinical studies (Bernard et al., 2014; Linhardt et al., 2022). First, 22 contextual questions are answered to adapt the rating system to a specific field of research (context). More specifically, these questions define the aspects that will be rated afterwards through the 34 design and implementation questions as well as what is considered as adequate or inadequate in that field of research (in this context). Note, that for the present review the answering of these questions was primarily based on the recently published consensus paper addressing the reporting and the experimental design of clinical and cognitive-behavioral neurofeedback studies (CRED-nf checklist; Ros et al., 2020). For detailed information about the customization performed for the current meta-analysis see Supplementary Material A. The DIAD algorithm delivers eight composite scores for each original study, which are finally summarized into the following four global scores of study bias: (A) *“fit between concepts and operations”* indicates whether the extent to which participants were handled, and outcomes were measured is in accordance with the definition of FMT-NF, (B) “*clarity of causal inference”* answers the question whether the study design allows an unequivocal conclusion regarding the effectiveness of the intervention, (C) “*generality of findings”* refers to the applied variation on participants, settings, and outcomes, and (D*) “precision of outcome estimation”* grades the quality and clarity of the reported effects.

### 2.3. Variables of Interest

To prove FMT-NF for its global effectiveness (primary aim of this study), it requires a reliable across-study comparison of changes in FMT as well as of behavioral changes induced by the neurofeedback intervention. To achieve this aim, five main contrasts of outcomes were initially defined: (1) The modulation of FMT amplitude during feedback, relative to a resting-state baseline (within session), (2) the modulation of FMT amplitude over time, relative to previous sessions (across sessions), (3) the changes of FMT amplitude from the resting state before to after a session (within rest), (4) the modulation of FMT amplitude from pre- to post-neurofeedback intervention during task performance (pre-post intervention) and (5) the modulation of task performance from pre- to post-neurofeedback (pre-post intervention behavior).

Because of the unexpected small number of original studies that finally met our inclusion criteria and could, thus could be included in each of the specified contrasts above, we had to deviate from our preregistration and adapted our investigation to the consideration of a broader range of reported effects and to the investigation of fewer contrasts. In detail, we finally defined effects of interest in the following way:

1. “Start-NF FMT modulation vs. end-NF FMT modulation” – (mod) comparing FMT amplitudes of the first and the last modulation run or between FMT amplitudes of the first and the last modulation session,
2. “Pre-NF FMT rest vs. post-NF FMT rest” – (bsl_pre_post) comparing FMT amplitudes during a resting-state measurement before and a resting-state measurement after a modulation run of a single session or comparing FMT amplitudes during resting state before and after the entire invention,
3. “Pre-NF FMT rest vs. during-NF FMT modulation” – (bsl_mod) comparing FMT amplitudes between resting-state baseline and modulation runs.

For each of these contrasts of interest, all available data were extracted from the original studies and included in a *global effect of FMT modulation* model (with the separate contrasts of interest 1-3 as random factors).

To better understand the influence of specific study parameters on the reported effects (secondary aim of the current study) additional variables were extracted. Specifically, we expected a) the number of neurofeedback sessions, b) the direction of the modulation, c) the use of individualized frequency bands as well as d) the use of several frequencies to induce moderating effects on all contrasts. However, due to the limited amount of available data, we did not conduct the moderator analysis for each FMT contrast specified above (1-3), but for the global FMT-NF effect model. Contrast-specific effects as well as the suggested potential moderators were assessed in subgroup analysis. A positive effect was always defined as a modulation in the intended direction, irrespective of the actual direction of the modulation, i.e., whether brain activity was up- or downregulated.

### 2.4. Effect Size Extraction

The definition of the term “modulation” and hence the reported effects in neurofeedback studies vary enormously (e.g., comparing amplitudes during modulation with amplitudes of the first session, or with amplitudes of the first resting state in that session, or with amplitudes of the resting state of the corresponding training day, or with amplitudes of the first run of a training session, etc.). To better disentangle this issue of nomenclature, we initially planned (see preregistration) to extract statistics (means, *Ms*, and standard deviations, *SDs*) of amplitudes corresponding to each of the three above specified contrasts (see section 2.3. Variables of Interest) from all original studies that met our inclusion criteria. This would have enabled us to calculate effects sizes ourselves and to conduct separate across-study comparisons for each contrast (1-3).

Since none of the identified 14 studies reported these statistics completely, all authors were contacted via email, but only one workgroup provided the requested data (Chen et al., 2022). Therefore, and in extension to our preregistered protocol, in eight cases (i.e., Brandmeyer and Delorme, 2020; Enriquez-Geppert et al., 2014b, 2014a; Eschmann et al., 2020; Reis et al., 2016; Rozengurt et al., 2016; Shtoots et al., 2021; Wang and Hsieh, 2013) effect sizes were calculated on the basis of the reported test statistics (*F*-values, *t*-values and *p*-values). In the remaining two cases (i.e., Rozengurt et al., 2017; Tseng et al., 2021), in which no test statistics of effects were reported in the text and we received no reply from the authors, effect sizes were extracted graphically using GetData Graph Digitizer (“GetData Graph Digitizer”, 2013). This procedure allowed us to retrieve six effect sizes for five of the original studies for contrast 1 (i.e., Brandmeyer and Delorme, 2020; Chen et al., 2022; Enriquez-Geppert et al., 2014a, 2014b; Reis et al., 2016), two effect sizes for contrast 2 (i.e., Chen et al., 2022; Reis et al., 2016) and nine effects sizes of seven studies for contrast 3 (i.e., Chen et al., 2022; Enriquez-Geppert et al., 2014b; Rozengurt et al., 2017, 2016; Shtoots et al., 2021; Tseng et al., 2021; Wang and Hsieh, 2013).

In total, 17 effects from 11 of the 14 studies included in this review could be extracted and were thus eligible for quantitative meta-analyses. All details on which data were available from each original study and the specific formulas used to calculate and convert the corresponding effect sizes are provided in supplementary material B.

### 2.5. Effect Size Calculation

Whenever means and standard deviations were available, either from provided data or retrieved visually from graphs in the original studies, the following formulas were applied to calculate Cohen’s *d* and the corresponding variance (*v_d_)* (Borenstein, 2009):

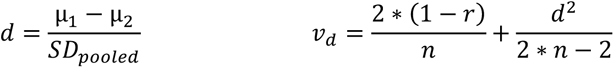

With *µ_1/2_* referring to the two means to be compared, *SD_pooled_* to the pooled standard deviation, *r* to the correlation between the two measurements and *n* to the sample size of the experimental group of the original study.

Whenever *F*-statistics were available, Cohen’s *d* and variances for effects comparing two groups were calculated using the following formulas (Borenstein, 2009; Thalheimer and Cook, 2002):

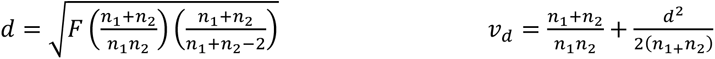

Where *F* corresponds to the *F*-statistic of an ANOVA and *n_1_* and *n_2_* to the sample sizes of the compared groups of the original studies.

If an effect size, such as Cohen’s *d* is calculated from a within-group effect, i.e., from a dependent *t*-test, the effect size usually suffers from a large overestimation (Dunlap et al., 1996). Hence, a corrected estimate accounting for an expected correlation (*r)* was used in these cases:

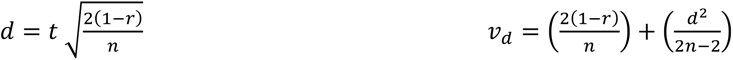

Where *t* corresponds to the *t-*statistic, *n* to the sample size and *r* to the estimated correlation between the two measurements in the original study. Since the therefore required correlation values were not reported, *r* was approximated. Specifically, we used the correlation values calculated from the one data set we gratefully received from Chen et al. (2022; correlations resting-state baseline (*M* = 1.13, *SD* = 0.57) vs. modulation (*M* = 1.88, *SD* = 0.30): *r* = .63; resting-state baseline vs. after neurofeedback (*M* = 1.20, *SD* = 0.53): *r* = .92; beginning (*M* = 1.96, *SD* = 0.38) vs. ending (*M* = 1.81, *SD* = 0.27) of neurofeedback (run 1 vs run 10): *r* = .67) as approximation. As an additional approximation and validity check, all effect sizes and all subsequent models were also calculated using a range of different *r-*values (most liberal: *r* = 0.74; most conservative: *r* = 0.97) based on previously reported retest-reliability of cognitive EEG (McEvoy et al., 2000). Respective results are briefly discussed but not presented in any detail here (available in supplementary material C, Fig. S2 and Fig S3).

Finally, as effect sizes have been shown to be prone to overestimation in small samples (Cumming, 2013), Hedges’ *g* correction was applied to all effect sizes and variances with the following formulas (Hedges, 1981):

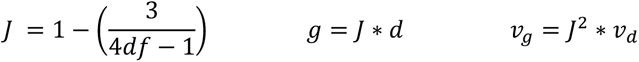

With *J* referring to the approximation of the correction factor and *df* to the degrees of freedom of the corresponding studies. The resultant values for Hedges’ *g* and *v_g_* represent the final input for all meta-analytical comparisons.

### 2.6. Meta-Analytic Synthesis of Results

The global effect of FMT-NF effectiveness was synthesized using all observed outcomes. Contrasts 1-3 and moderator analyses were performed in the same manner as the analysis of the global effect, with only the respective data included in these models.

In common meta-analytic models, it is assumed that the sampling errors of observed effects are independent. This rule is violated, if several observed effect sizes are based on the same sample or on partially overlapping samples; in those cases, the effects (or more specifically their sampling errors) are dependent. Thus, in all cases in which we included dependent effects (i.e., the global meta-analytic model, contrast 3 and all moderator analysis), the variance-covariance matrix (*V*) of the sampling errors was approximated using the *metafor vcalc()* function. *V* was then used as input into the metafor function *rma.mv()*, fitting the multivariate random-effects model (REM) to the data, while for contrast 1 and 2, the *rma.uni()* function for the univariate model was used. To account for the random effects of each study and for the previously defined contrast (1-3; see section 2.3. Variables of Interest), the input parameter *random* was set to *∼ 1 | study_id/subeffect* for the global model and all moderator analysis. For the three contrast-specific analysis *random* was set to *∼ 1 | study_id*.

Note that in cases of dependent effects (contrast 3 and the moderator analysis) cluster-robust estimates of the variance-covariance matrix were obtained for all multivariate REMs, by subjecting the model to the *robust()* method with the improved inference methods of the *clubSandwich* package in a last step. This was necessary in those cases, as *V* represents an approximation, and the dependencies might not be fully accounted for by the random effects structure.

Study heterogeneity was assessed with *Q*-tests (Cochran, 1954), independently among the studies included in each model. The respective results are reported for each model in a forest plot by *Q*-, *p- and I*²-values. In univariate models, variance is described by =τ^2^, while the two variance components in multivariate models are indicated by σ²_study_ (between-cluster heterogeneity; the studies) and σ²_effect_ (within-cluster heterogeneity; the subeffects 1-3 within studies). Both components were visually investigated by profile likelihood plots. Further, to index the dispersion of effects, a prediction interval was calculated for the observed outcomes and included in the forest plot (Riley et al., 2011).

Outliers within the effects of original studies were defined in accordance to Viechtbauer and Cheung (2010) as effects with a studentized residual larger than 1.96. Additionally and as also recommended by Viechtbauer and Cheung (2010), influential effects, i.e., effects whose observed outcomes strongly influence the model outcome, were defined when Cook’s distance was larger than the median plus six times the interquartile range. To allow for most comprehensive insights into the robustness of across-study effects, all meta-analytical effects are reported with and without the inclusion of effects that fulfill both criteria. We refer to these effects as “influential outliers” in the following.

The programming language *R* (version 4.2.2, R Core Team, 2020) and the *metafor* package (version 3.8.1) were used to carry out all meta-analyses and to generate all plots (Viechtbauer, 2010). Statistical significance was accepted at *p* < .05.

### 2.7. Publication Bias

Publication bias was assessed graphically with a funnel plot. Additionally, funnel plot asymmetry was investigated with rank-correlation tests (Begg and Mazumdar, 1994) and file-drawer analyses. File-drawer analyses were conducted following the approach of Rosenthal (Failsafe-*N,* 1979), i.e., by identifying the number of null results necessary to reduce the *p*-value to an alpha level < .05, as well as following the approach of Orwin (1983), i.e., by identifying the number of null results necessary to reduce the average outcome to an effect below “clinical” significance; here we choose a small effect of Hedges’ *g* = 0.2.

### 2.8. Data and Code Availability

All software and packages used in the current study are open source. The code used to conduct the meta-analyses and to create corresponding figures included in this paper is available in a GitHub repository (https://github.com/iamraP/MetaAnalysis/). This repository further includes code to create the plots for each contrast-specific analysis (also provided in supplementary C) as well as the data extracted from the original studies (see also Supplementary Material B).

## 3. Results

### 3.1. Study Quality – Study DIAD

The global DIAD quality scores of each study are listed in Fig. 2 on the right side next to the forest plot. More detailed results of the design and implementation questions, the contextual questions as well as the rating algorithm are provided in Supplementary Material A.

**Fig. 2.**
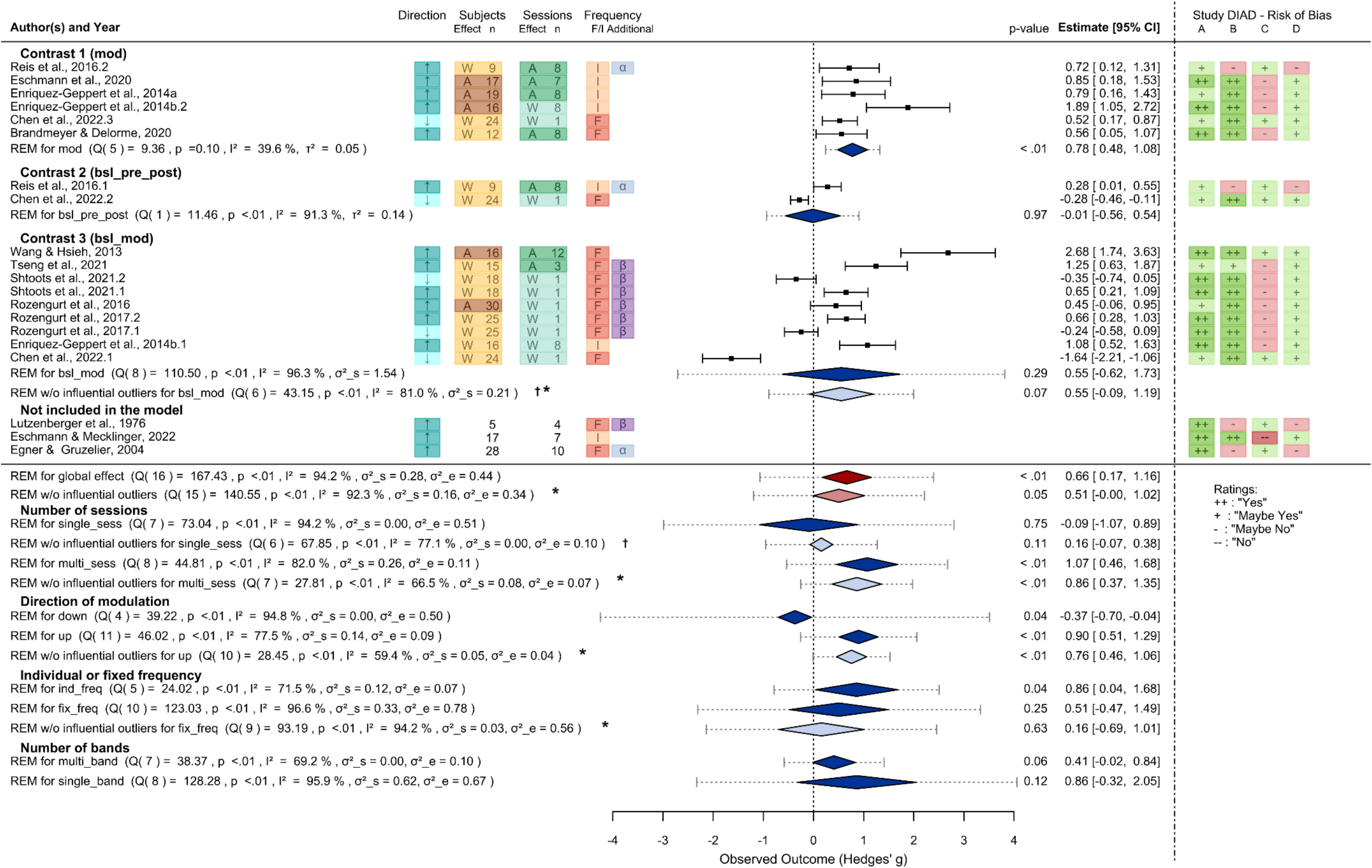
Forest Plot of the multivariate random effect model (REM) including all studies with the contrast-specific analysis depicted below. Effects are identified by author, year, and an additional number to distinguish them (whenever several effects were obtained within a single study). Study characteristics with importance to the moderator analyses are displayed in the following way: Direction of modulation: **up (↑) / down (↓)**; number **(n)** of **subjects** in the experimental group and **sessions**; whether the **effect** was measured **within (W)** and **across (A) subjects** or **sessions**; whether a **fixed (F)** or **individual (I) frequency** band was used and if the modulation of an **additional frequency** band was intended. The contrasts of interest (1-3) correspond to the extracted effects: **“mod”** - Start-NF FMT modulation vs. end-NF FMT modulation, **“bsl_pre_post” -** Pre-NF FMT rest vs. post-NF FMT rest and “**bsl_mod**” - Pre-NF FMT rest vs. during-NF FMT modulation. The observed outcomes of FMT modulation are presented as the standardized mean differences (Hedges’ g). Means of individual effects are depicted by black squares with solid whiskers which indicate the 95%-Confidence Interval (CI). Diamonds with dotted whiskers depict the 95%-CI of the REMs for the global effects (red), the contrast-specific and moderator effects (blue) and their respective prediction intervals (dotted whiskers). Lighter colours indicate the exclusion of influential outliers. These excluded studies (Chen et al., 2022.1 (**†**) and Wang & Hsieh, 2013(**\***)) are identified in the corresponding columns. Study heterogeneity is reported by **I²**. While for univariate models the between-study variance is given by **τ²**, for multivariate models the between study-variance is indicated by **σ²_s** and the within-study variance by **σ²_e**. For each reported effect the results of the risk of bias assessment of the corresponding study are presented in the four most right columns: (**A**) Fit between Concept & Operations, (**B**) Clarity of Causal Inference, (**C**) Generalizability of Findings and (**D**) Precision of Outcome Estimation.

*Fit between concepts and operations:* All of the 14 studies included in the review demonstrated sufficient “fit between concepts and operations” (Yes: 7, Maybe Yes: 7), which is most likely due to our strict exclusion criteria, assuring that all studies were implemented in a way that outcome measures were consistent with the definition of FMT neurofeedback. Detected flaws that led to a lower rating in this DIAD category (Maybe Yes instead of Yes) refer in all cases to the question of whether all information is provided to replicate the study (question 1.1.2). Especially the specification of the instructions given to participants were insufficiently described.

*Clarity of the causal inference:* Comparably good results were obtained for “clarity of the causal inference” (Yes: 10, Maybe Yes: 1, Maybe No: 3). Only one study reported differential attrition between experimental groups and only two studies suffered from severe (>10%) overall attrition. Less positive ratings in this DIAD category (Maybe Yes or Maybe No) were either due to the recruitment from the same local pool (question 2.2.2) or to non-randomized allocation of participants (question 2.2.1).

*Generalizability of finding:* Due to the lack of heterogeneous study populations (question 3.2.1) and variations in experimental settings (question 3.2.4 and 3.2.5), the DIAD scores for the category “generalizability of finding” were overall rather low (Maybe Yes: 5, Maybe No: 8, No: 1).

*Precision of outcome estimation:* The low sample sizes inherent to all studies directly affected the precision of outcomes (question 4.1.3). Therefore, similarly low scores were obtained in this domain of the DIAD by all studies (Maybe Yes: 11, Maybe No: 3). Also, the reporting of outcomes (question 4.2.3) was overall rated as insufficient. Means and standard deviations were only retrievable from graphics and effect sizes were mostly not reported.

### 3.2. Systematic Review

#### 3.2.1. Study Characteristics

Table 1 presents a detailed overview of design characteristics of the 14 studies included in the qualitative review. Average sample sizes for experimental and control groups were 17 (*SD*: ± 6.9) and 14.8 (*SD*: ± 5.8) participants per group, respectively. The durations of neurofeedback-interventions varied between one day with only one neurofeedback session (i.e., Chen et al., 2022; Rozengurt et al., 2017, 2016; Shtoots et al., 2021) and four weeks including 12 sessions (i.e., Wang and Hsieh, 2013). On average, multi-session neurofeedback-intervention studies comprised 7.5 sessions (*SD*: ± 2.6) (i.e., Brandmeyer and Delorme, 2020; Egner and Gruzelier, 2004; Enriquez-Geppert et al., 2014a, 2014b; Eschmann et al., 2020; Eschmann and Mecklinger, 2022; Lutzenberger et al., 1976; Reis et al., 2016; Tseng et al., 2021; Wang and Hsieh, 2013).

Five of the 14 studies used individual frequency training (IFT). Here, the experimenter adjusts the frequency band that is modulated during the neurofeedback training to peaks in the power spectrum of individual participants instead of using the same fixed frequency band for all participants (i.e., Enriquez-Geppert et al., 2014b, 2014a; Eschmann et al., 2020; Eschmann and Mecklinger, 2022; Reis et al., 2016). Eight of the 14 included studies only provided feedback of the theta frequency band (i.e., Brandmeyer and Delorme, 2020; Chen et al., 2022; Enriquez-Geppert et al., 2014a, 2014b; Eschmann et al., 2020; Eschmann and Mecklinger, 2022; Lutzenberger et al., 1976; Wang and Hsieh, 2013), while the remaining six aimed at the simultaneous modulation of other frequency bands (i.e., beta: Rozengurt et al., 2017, 2016; Shtoots et al., 2021; Tseng et al., 2021; alpha: Egner and Gruzelier, 2004; Reis et al., 2016). In two FMT neurofeedback studies the neurofeedback was preceded by another biofeedback intervention: One study used real and sham feedback to modulate heart rate and frontal muscle tension (Lutzenberger et al., 1976), while another study used neurofeedback to modulate frontal alpha brain activity (Reis et al., 2016).

Eleven out the 14 considered investigations focused exclusively on the upregulation of FMT (i.e., Brandmeyer and Delorme, 2020; Egner and Gruzelier, 2004; Enriquez-Geppert et al., 2014a, 2014b; Eschmann et al., 2020; Eschmann and Mecklinger, 2022; Lutzenberger et al., 1976; Reis et al., 2016; Rozengurt et al., 2016; Tseng et al., 2021; Wang and Hsieh, 2013), one study strived only for downregulation of FMT (Chen et al., 2022) and two studies intended to investigate modulations of FMT in both directions, i.e., using one direction of the modulation as control condition (i.e., Rozengurt et al., 2017; Shtoots et al., 2021).

The most frequently implemented control condition in the included studies was sham-neurofeedback (i.e., Brandmeyer and Delorme, 2020; Chen et al., 2022; Enriquez-Geppert et al., 2014a, 2014b; Eschmann et al., 2020; Eschmann and Mecklinger, 2022; Lutzenberger et al., 1976; Reis et al., 2016), in which participants receive feedback from another participant or from a changing, randomly selected frequency band. Further, the modulation of another fixed frequency band (i.e., beta: Rozengurt et al., 2016), other electrode positions (Egner and Gruzelier, 2004) or diverging instructions (Chen et al., 2022) were used as control. Four studies used control groups not receiving any neurofeedback, but performing movie-watching (i.e., Rozengurt et al., 2017), cognitive tasks (i.e., Reis et al., 2016), or not participating in any intervention (i.e., Shtoots et al., 2021; Tseng et al., 2021).

The feedback itself was mostly provided visually (i.e., Brandmeyer and Delorme, 2020; Enriquez-Geppert et al., 2014a, 2014b; Eschmann et al., 2020; Eschmann and Mecklinger, 2022; Reis et al., 2016; Rozengurt et al., 2017, 2016; Shtoots et al., 2021), or auditorily (i.e., Chen et al., 2022; Egner and Gruzelier, 2004; Lutzenberger et al., 1976; Tseng et al., 2021), while combined audio-visual feedback was delivered only in one study (Wang and Hsieh, 2013). In ten of the 14 included studies, this feedback was derived from the EEG-signal measured at a single electrode (Fz; i.e., Brandmeyer and Delorme, 2020; Chen et al., 2022; Eschmann et al., 2020; Eschmann and Mecklinger, 2022; Lutzenberger et al., 1976; Reis et al., 2016; Rozengurt et al., 2017; Shtoots et al., 2021; Tseng et al., 2021; Wang and Hsieh, 2013), two investigations determined the feedback on the basis of activity measured from multiple but only frontal electrodes (i.e., Enriquez-Geppert et al., 2014b, 2014a) and two studies also considered parietal electrodes, either for a subset of the participants in the experimental group (i.e., Rozengurt et al., 2016) or as a control condition (i.e., Egner and Gruzelier, 2004).

Not all included studies aimed at influencing a specific behavior. Two studies focused on the feasibility of frontal theta modulation itself (i.e., Egner and Gruzelier, 2004; Lutzenberger et al., 1976), while other studies examined whether the neurofeedback intervention could lead to improvement in the domains of memory and cognitive performance (i.e., Enriquez-Geppert et al., 2014b, 2014a; Eschmann et al., 2020; Eschmann and Mecklinger, 2022; Reis et al., 2016; Rozengurt et al., 2017; Shtoots et al., 2021; Tseng et al., 2021; Wang and Hsieh, 2013), motor learning (i.e., Rozengurt et al., 2016) and sports performance (i.e., Chen et al., 2022), or meditation and mind wandering (i.e., Brandmeyer and Delorme, 2020). Finally, Chen et al. (2022) applied different types of instructions to test, if a particular strategy on how to modulate brain activity helps or hinders participants in their learning.

Overall, nine of the 14 included studies instructed participants explicitly to use a specific strategy for the modulation. Instructed strategies to increase FMT activity include meditation (i.e, Brandmeyer and Delorme, 2020), relaxation (i.e.,Egner and Gruzelier, 2004; Rozengurt et al., 2016), emotions and memories (i.e.,Enriquez-Geppert et al., 2014a), mental and motor imagery (i.e.,Enriquez-Geppert et al., 2014a; Eschmann et al., 2020; Eschmann and Mecklinger, 2022), arithmetic operations (i.e., Enriquez-Geppert et al., 2014a; Eschmann et al., 2020; Eschmann and Mecklinger, 2022; Tseng et al., 2021), and others (i.e., Enriquez-Geppert et al., 2014b; Tseng et al., 2021). The only study which provided instructions for FMT downregulation asked their participants “to decrease conscious effort“ (Chen et al., 2022). Four studies refrained from using any specific instructions on how to modulate, in order to allow participants to solely focus on the feedback signal, and to develop their own strategies (i.e., Lutzenberger et al., 1976; Rozengurt et al., 2017; Shtoots et al., 2021; Wang and Hsieh, 2013). In the remaining report the instruction process of the participants was not described at all (Reis et al., 2016).

#### 3.2.2. Modulation Within Neurofeedback Session(s)

Eight of the 14 reports included in the qualitative review described the modulation of FMT within sessions (i.e., Chen et al., 2022; Egner and Gruzelier, 2004; Enriquez-Geppert et al., 2014b; Eschmann and Mecklinger, 2022; Lutzenberger et al., 1976; Rozengurt et al., 2017, 2016; Shtoots et al., 2021). Only one of these studies reported a responder rate for the within session modulation: 76%, based on a theta increase of > 5% relative to a resting-state baseline (i.e., Rozengurt et al., 2017).

Upregulation of FMT via neurofeedback was in most studies successful (i.e., Enriquez-Geppert et al., 2014b; Eschmann and Mecklinger, 2022; Rozengurt et al., 2017, 2016; Shtoots et al., 2021). However, Reis et al. (2016) only observed significant FMT upregulation within sessions when examining FMT power relative to the broadband of 0.2 – 35 Hz (the relative power describes which proportion of the measured EEG is FMT), but not in absolute values and Egner and Gruzelier (2004) as well as Lutzenberger et al. (1976) could not detect any within session effects for modulation runs. However, Lutzenberger et al. (1976) noted a significant decrease of theta when considering transfer runs, in which participants were asked to increase theta but did not receive feedback.

More ambiguous results were observed for the downregulation of FMT via neurofeedback. Aiming for an increase in the beta/theta-ratio, Rozengurt et al. (2017) reported only a slight, non-significant decrease, while Shtoots et al. (2021) described an increase of theta even within their single session protocol. Chen et al. (2022) observed significant reduction during the modulation runs, but only for participants receiving function-specific instructions (i.e., paying attention and decreasing conscious effort).

#### 3.2.3. Modulation Across Neurofeedback Sessions

All of the ten studies which implemented a multi-session protocol provided information about potential modulation changes across sessions. Two of these studies also reported responder rates for across session modulations, both of these studies aimed for theta upregulation, and both of them observed successful modulation in 75% of their participants (i.e., Brandmeyer and Delorme, 2020; Enriquez-Geppert et al., 2014b). However, Brandmeyer and Delorme (2020) defined non-responders as participants whose daily neurofeedback scores differed with three or more *SD*s from the mean neurofeedback score of all participants, while Enriquez-Geppert et al. (2014b) defined non-responders as participants that did not show any up-regulation relative to the first session.

All ten multi-session studies focused on upregulating FMT, while no single study strived for downregulation. Two of these ten studies did not detect any significant change across sessions (i.e., Egner and Gruzelier, 2004; Tseng et al., 2021), while eight investigations observed increased FMT over time (i.e., Brandmeyer and Delorme, 2020; Enriquez-Geppert et al., 2014a, 2014b; Eschmann et al., 2020; Eschmann and Mecklinger, 2022; Lutzenberger et al., 1976; Reis et al., 2016; Wang and Hsieh, 2013).

#### 3.2.4. Effects of FMT Neurofeedback on Resting-State or Task-Related EEG

Four of the 14 reviewed studies investigated the effects of FMT modulation via neurofeedback on resting-state EEG. Two of them reported significant changes after the neurofeedback session in the same direction as the modulation, i.e., FMT increase after upregulation (Reis et al., 2016; Wang and Hsieh, 2013). In contrast, Chen et al. (2022) could not detect any effect after FMT downregulation and Lutzenberger et al. (1976) reported a decrease of FMT after intended upregulation when comparing baselines before and after the neurofeedback within a session, but an increase when comparing FMT across sessions.

Six of the 14 included studies investigated the effects of FMT neurofeedback on EEG acquired during tasks. Chen et al. (2022) observed that only the group receiving function-specific instructions before neurofeedback training demonstrated decreased FMT during golfing puts, while the decrease aligned with the direction of the NF training. Contrarily, Eschmann and colleagues (2020, 2022) investigated event-related synchronization and reported effects in the opposite direction of their FMT upregulation during certain tasks. More specifically, they detected a non-significant trend of reduced pre-stimulus FMT during a source memory task (Eschmann et al., 2020), as well as a trend for decreased FMT post-stimulus in the retention condition of a Delayed-Match-To-Sample (DMTS) task (Eschmann and Mecklinger, 2022). Notably, the effect for the retention condition was not present immediately after the last neurofeedback session, but 13 days later. Increased task-related FMT was also found after training upregulation during a 3-back task, a task-switching paradigm, a Stop-Signal task and a Stroop task (Enriquez-Geppert et al., 2014a), as well as during motor sequence learning (Rozengurt et al., 2016). Finally, Lutzenberger et al. (1976) demonstrated that FMT decreased during aversive movie watching in the sham neurofeedback control condition, while it remained stable for the group which learned FMT upregulation.

#### 3.2.5. Modulation of Behavior after FMT Neurofeedback

The majority of studies assessing potential behavioral changes induced by FMT neurofeedback belong to the domain of cognitive control. This refers to seven out of the 12 studies, which tested for behavioral changes in various cognitive functions. However, while effective behavioral modulation could be observed in respect to some of these functions, behavioral performance in other functions failed at effective modulation. For example, inhibitory control could not be improved if operationalized by behavioral performance in the Stroop task (i.e., Enriquez-Geppert et al., 2014a; Eschmann and Mecklinger, 2022), the Stop-Signal task (i.e., Enriquez-Geppert et al., 2014a), the Sustained-Attention-To-Response task (i.e., Brandmeyer and Delorme, 2020) nor by performance in the Local-Global task (i.e., Brandmeyer and Delorme, 2020). In contrast, Enriquez-Geppert et al. revealed improved task-switching after FMT upregulation, arguing that memory updating and mental set shifting rely on proactive instead of reactive control, as it would be the case for motor inhibition and conflict monitoring (Enriquez-Geppert et al., 2014a).

Eschmann and Mecklinger (2022) observed no enhanced performance compared to a control group in a Delayed-Match-To-Sample Manipulation task involving mirroring (and thus remembering) of the stimulus directly and 13 days after FMT upregulation. However, they demonstrated a significantly superior performance of the neurofeedback-group in a regular Delayed-Match-To-Sample task, but only 13 days after the last training session. Notably, theta changes were associated with better performance in both tasks.

Successfully enhanced working memory performance after FMT upregulation has been investigated in seven studies and was demonstrated for the n-back task (i.e., Brandmeyer and Delorme, 2020; Enriquez-Geppert et al., 2014a), the Sternberg task (i.e., Wang and Hsieh, 2013), a matrix rotation task (i.e., Reis et al., 2016) and the Attention Network task (i.e., Wang and Hsieh, 2013). Notably, Reis et al. (2016) even revealed positive correlations between the theta gradient during neurofeedback and the amount of change in accuracy (task performance) during matrix rotation, thus providing promising hints for the specificity of that intervention. In the Attention Network task, Wang and Hsieh (2013) observed improvement in the conflict network, i.e., less slowing in incongruent trials as compared to congruent ones, but not in the alerting network, i.e., sensitivity to cues. Improvements in the orientation network were only present in an elderly (age: 65 ± 3.3 years), but not in a younger subsample (age: 22.6 ± 1.6 years). However, no improvements in working memory performance were reported by Tseng et al. (2021) after FMT upregulation in a Backward Digit Span task.

Further, FMT upregulation has been shown to correlate with increased behavioral performance in tasks assessing semantic memory (i.e., Tseng et al., 2021), source memory (i.e., Eschmann et al., 2020) and visual-spatial memory (i.e., Shtoots et al., 2021). In contrast, no association could be observed between FMT upregulation and visual-spatial memory improvements by Eschmann et al. (2020). However, they revealed a positive correlation between increased performance and the amount of FMT upregulation induced by neurofeedback, showing that roughly 30% of the enhancement in the neurofeedback-group could be explained by the FMT change during neurofeedback (i.e., Eschmann et al. 2020). Similarly, Tseng et al. (2021) detected that the forgetting rate in the semantic memory task was inversely related to the difference of FMT power between one and three days after neurofeedback training.

Concerning item memory, Eschmann et al. (2020) could not observe any alterations after FMT upregulation via neurofeedback, while Rozengurt et al. (2017) and Tseng et al. (2021) observed significant enhancements in this domain. Furthermore, Rozengurt and colleagues (2017) showed that inverse modulation (downregulation of FMT) was associated with memory decline and that the percentage of theta change during neurofeedback correlated significantly with the amount of successfully recalled objects immediately after the session.

Finally, a positive correlation between FMT upregulation and enhanced motor learning was identified in a Finger-Tapping task by Rozengurt et al. (2016). Aiming at FMT reduction, Chen et al. (2022) showed enhanced performance during golf putting, but only in a group receiving function-specific instructions instead of solely being asked to modulate the feedback signal. No correlation between FMT-modulation and performance changes was found.

Overall, studies aiming for the modulation of memory performance appeared to be more successful than studies seeking for improving inhibitory control or interference suppression. In total, 21 different cognitive tasks were employed in the ten included studies which investigated cognitive performance. Positive correlations were reported for the Delayed-Match-To-Sample task (i.e., Eschmann and Mecklinger, 2022), matrix rotation (i.e., Reis et al., 2016), semantic memory (Tseng et al., 2021), source memory (Eschmann et al., 2020) and item memory (i.e., Rozengurt et al., 2017). Contradicting results were obtained for visual spatial memory (i.e., Eschmann et al., 2020; Shtoots et al., 2021). Sports (golfing) performance and motor sequence learning were only explored in one study each, in which correlational evidence for modulation was only detected for FMT increase and motor sequence learning (i.e., Rozengurt et al., 2016).

### 3.3. Meta-Analytic Synthesis

Seventeen effects of 11 different studies were eligible for inclusion in quantitative meta-analyses. In total, 189 participants were assigned to experimental groups (FMT modulation), while control groups comprised 255 subjects. Ten of those studies reported effects for upregulation (i.e., Brandmeyer and Delorme, 2020; Enriquez-Geppert et al., 2014a, 2014b; Eschmann et al., 2020; Eschmann and Mecklinger, 2022; Reis et al., 2016; Rozengurt et al., 2017, 2016; Shtoots et al., 2021; Tseng et al., 2021; Wang and Hsieh, 2013), while three investigations demonstrated effects for downregulation (i.e., Chen et al., 2022; Rozengurt et al., 2017; Shtoots et al., 2021). Of the 17 effects, seven focused on neurofeedback-induced modulation of frontal midline theta across different sessions of an intervention (i.e., Brandmeyer and Delorme, 2020; Enriquez-Geppert et al., 2014a; Eschmann et al., 2020; Reis et al., 2016; Tseng et al., 2021; Wang and Hsieh, 2013), while ten effects reflect modulation changes within sessions (i.e., Chen et al., 2022; Enriquez-Geppert et al., 2014b; Rozengurt et al., 2017, 2016; Shtoots et al., 2021). Further, 12 of the 17 included effects were measured within the same group of participants (i.e., Brandmeyer and Delorme, 2020; Chen et al., 2022; Enriquez-Geppert et al., 2014b; Reis et al., 2016; Rozengurt et al., 2017; Shtoots et al., 2021; Tseng et al., 2021), while five effects were calculated based on the comparison with a control group (i.e., Enriquez-Geppert et al., 2014b, 2014a; Eschmann et al., 2020; Rozengurt et al., 2016; Wang and Hsieh, 2013).

Which effects were based on within vs. across session modulation and which on within vs. across subject measurements, as well as other important study parameters, are listed in the corresponding rows and columns of the forest plot in Fig. 2. To differentiate between multiple effects obtained in a single study, those effects were labeled with an additional number following the year of the publication (e.g., Chen et al.2022.1, Chen et al.2022.2).

#### 3.3.1. Outliers and Influential Studies

In the context of the global meta-analytical model, two effects showed studentized residuals ± 2.21 (Chen et al. 2022.1a; Wang and Hsieh 2013), but only one effect (Wang and Hsieh, 2013) also fulfilled the criteria of Cook’s distance > 1.96 and is, therefore, considered as influential outlier. With respect to the subgroup analyses, two effects fulfill both criteria (Chen et al., 2022.1, Wang and Hsieh, 2013) and were, thus, treated as influential outliers in five of the 11 planned subgroup analyses (see Fig. 2 for details).

#### 3.3.2. Global Effect on Frontal Midline Theta Induced by Neurofeedback Interventions

Subsuming over all 17 effects reflecting neurofeedback-induced modulation of frontal midline theta in healthy adult participants, the multivariate random effects model indicated a significant positive cross-study effect of medium size (*p* = .026, Hedges’ *g* = .66; 95%-CI [−0.62, 1.73]; Fig. 2). When excluding the two effects defined as influential outliers (Chen et al. 2022.1; Wang and Hsieh 2013), this cross-study effect no longer reached statistical significance (*p* = .504, Hedges’ *g* = .51; 95%-CI [−0.00, 1.02]). In both cases, significant across-study heterogeneity in reported effect sizes was observed (Q(16) = 167.43, *p* < .0001 and Q(15) = 140.55, *p* < .0001, respectively).

#### 3.3.3. Subanalyses

To reveal more detailed insights into the global meta-analytical effect, a total of 11 subanalyses evaluating across-study effect sizes for smaller selections of studies were performed. Specifically, we compared effects resulting from the three different contrasts considering different measurement time points (see section 2. Variables of Interest) as well as the effects of different study parameters, i.e., single- and multi-session protocols, up- and downregulation of FMT, individual and fixed frequency bands, and modulation of multiple bands or only one. All subanalyses results are included in Fig. 2, while independent forest plots for all subanalyses are presented in the Supplementary Material C Fig. S1a-S1p.

Five of the 11 investigated effects, with seven models in total, turned out to be significant.

##### 3.3.3.1. Contrasts of Interest (1-3)

First of all, the univariate random effects model for contrast 1, i.e., comparing the beginning and ending of a neurofeedback session or intervention (mod), delivered a significant effect of medium size (*p* < .01, Hedges’ *g* = .78; 95%-CI [0.48, 1.08]) and a moderate amount of across-study heterogeneity in reported effect sizes with a non-significant Q-test (Q(5) = 9.36, *p* = .10, I² = 39.6 %, = *τ*² = 0.05). Thus, it can be assumed that the observed across-study effects belong to the same underlying true effect and the dispersion is caused by sampling error. For contrasts 2 (bsl_pre _post) and 3 (bsl_mod) no significant effects were found (bsl_pre_post: *p* = .97, Hedges’ *g* = -.01, 95%-CI = [-0.56, 0.54]; bsl_mod: *p* = .29, Hedges’ *g =* .55, 95%-CI = [-0.62, 1.73] with significant across-study heterogeneity (bsl_pre_post: Q(1) = 11.46, *p* < .01, I² = 91.3 %, *τ²* = 0.14; bsl_mod: (Q(8) = 110.50, *p* < .01, I² = 96.3 %, σ²_study_ = 1.54).

For contrast 1 (mod) and 2 (bsl_pre_post) no influential outliers were detected, while for contrast 3 (bsl-mod) two influential outlier studies were identified (Wang and Hsieh, 2013 and Chen et al. 2022.1). Removing these two studies from the models reduced study heterogeneity significantly (Q(6) = 43.15, *p* < .01, I² = 81.0 %, σ²_study_ = 0.21), but did not affect the effect-size model itself (*p* = .07, Hedges’ *g* = .55, 95%-CI [-0.09, 1.19]). Nonetheless, the large overlap of the confidence intervals of the four models of the three contrasts suggests the absence of significant differences between across-study effects.

##### 3.3.3.2. Number of Sessions

Secondly, the multivariate random effects models for the moderator analysis on multi-session protocols, with and without the influential outlier (Chen et al. 2022.1), reached statistical significance and provided a large cross-study effect (with influential outliers: *p* < .01, Hedges’ *g* = 1.07, 95%-CI [0.46, 1.68]; without the influential outlier: *p* < .01, Hedges’ *g* = 0.86, 95%-CI [0.37, 1.35]). A wide dispersion of the original study effect sizes was obvious and high I²-values suggest that a large proportion of the variance between studies is real and most likely not attributable to sampling error (Q(8) = 44.81, *p* > .01, I² = 82.0 %, σ²_study_ = 0.26, σ²_effect_ = 0.11; Q(7) = 27.81, *p* < 01, I² = 66.5 %, σ²_study_ = 0.08, σ²_effect_ = 0.07). The models for single-session protocols were not significant neither with, nor without the influential outlier (Wang and Hsieh, 2013) – detailed model characteristics are reported in Fig. 2. Differently estimated effects are indicated by the barely overlapping confidence intervals of the models for single- and multi-session protocols without the influential outliers.

##### 3.3.3.3. Direction of Modulation

Thirdly, the multivariate random effects model testing FMT upregulation protocols showed a large and significant effect in the original model (*p* < .01, Hedges’ *g* = 0.90, 95%-CI [0.51, 1.29]), which was reduced to an effect of medium size after one study identified as an influential outlier (Wang and Hsieh, 2013) was excluded (*p* < .01, Hedges’ *g* = 0.76, 95%-CI [0.46, 1.06]). Significant across-study heterogeneity in effect sizes was detected in both models, and, as indicated by I², less than 25% of the dispersion of effect sizes can be attributed to sampling error in the original model (Q(11) = 46.02, *p* < .01, I² = 77.5 %, σ²_study_ = 0.14, σ²_effect_ = 0.09). The exclusion of the influential outlier increased this to roughly 40% (Q(10) = 28.45, *p* < .01, I² = 59.4 %, σ²_study_ = 0.05, σ²_effect_ = 0.04).

Fourth, the model testing for the FMT downregulation reached statistical significance with a negative across-study effect of medium size (*p* = .04, Hedges’ *g* = -0.37 [-0.70, -0.04]). No influential outlier studies were detected in this model. The observed across-study heterogeneity in effect sizes was significant and extremely high (Q(4) = 39.22, *p* < .01, I² = 94.8 %, σ² _study_ = 0.00, σ²_effect_ = 0.50). Importantly, the calculation of this across-study effect was only based on five effect sizes with three effect sizes stemming from a single dataset. Further, a large overlap between this subgroup and the subgroup of effects extracted from single-session protocols was present. Notably, this was the only other model with an – although not significant, but still overall – negative effect size estimate (*p* = .75, Hedges’ *g* = -0.09, 95%-CI [-1.07, 0.89]). Finally, the absence of overlap in confidence intervals of the up- and downregulation models again suggests differentiable underlying effects.

##### 3.3.3.4. Individual versus Fixed Frequency

Fifth, a large and significant effect was observed for studies using individual frequencies for neurofeedback-training (*p* = .04, Hedges’ *g* = 0.86, 95%-CI [0.04, 1.68] with significant across-study heterogeneity (Q(5) = 24.02, *p* < .01, I² = 71.5 %, σ²_study_ = 0.12, σ²_effect_ = 0.07). No influential outlier was detected in the context of this model. In contrast, no significant effect was found for the models for fixed-frequency protocols, irrespective of whether the influential outlier (Wang and Hsieh, 2013) was included or not. Note that a large overlap between both confidence intervals does not support the assumption that the observed effects for individual- and fixed-frequency trainings belong to separate distributions.

##### 3.3.3.5. Number of Bands

The moderator analyses for multi- and single-band protocols yielded no significant effects and no influential outliers were detected in the context of these two models. Further, the nearly complete overlap of the confidence intervals suggests the absence of significant difference between the models (Fig. 2).

Overall, it is important to note that all 18 submodels demonstrated significant (*p* < .01) across-study heterogeneity in the reported effects. All but four comparisons resulted in I²-values above 70% accompanied by wide dispersions of their prediction intervals. This suggests different underlying across-study effects, or more specifically, that different parameter choices in study design significantly influence the size of the observed effects.

#### 3.3.4. Publication Bias

Visual inspection of the funnel plot (see Fig. 3), as well as the significant results of the rank correlation test (Kendall’s τ= 0.47, *p* < .01), assessing the association between effect sizes and standard errors, indicate significant funnel plot asymmetry. This implies that the higher the effect reported in a given study, the larger the error associated with this effect, and ultimately suggests the presence of publication bias. Further support for this assumption comes from the file-drawer analysis (Rosenthal’s fail-safe N), which was also significant (*p* < .0001, failsafe-N = 282) and Orwins approach indicated that 33 null-results would be required to reduce the effect to Hedges’ *g* = 0.2. Note, however, that these effects might also be introduced by the highly heterogeneous study designs and the limited number of studies employed.

**Fig. 3.**
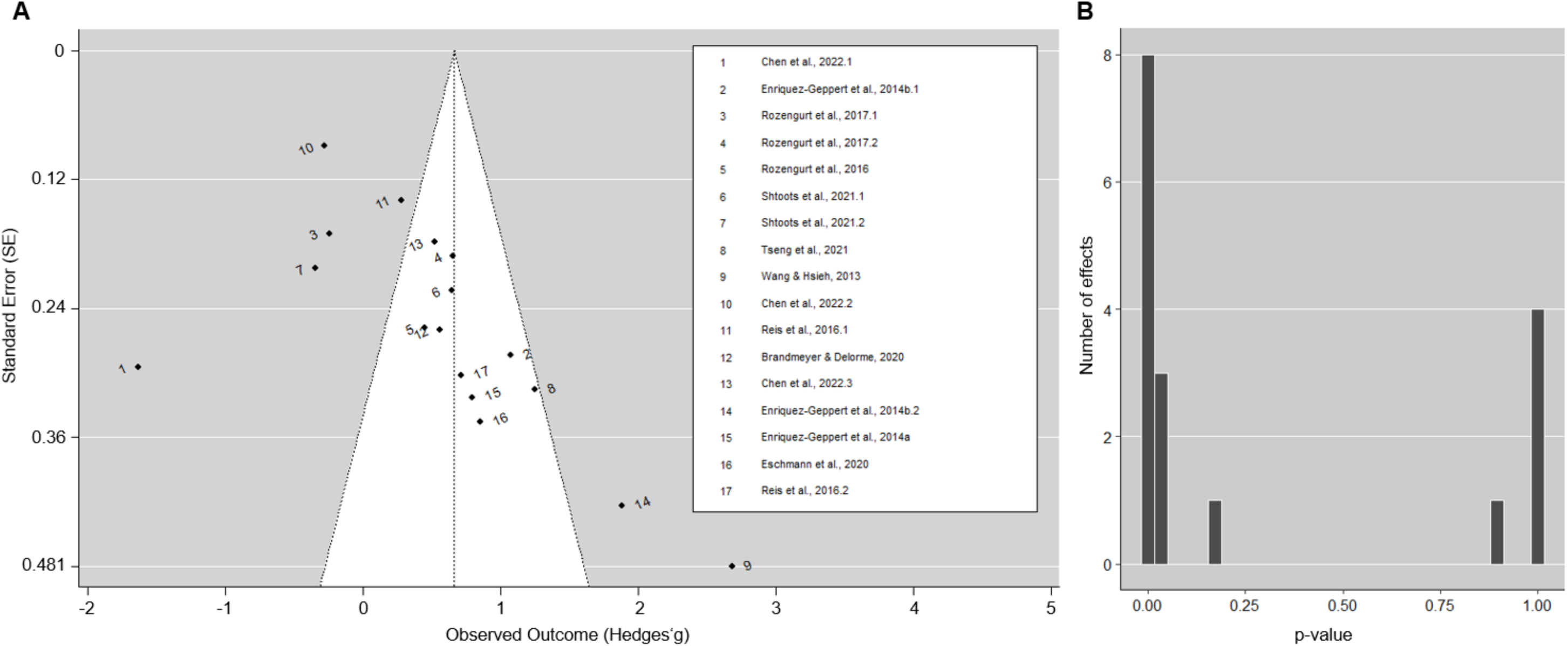
Publication bias assessment. **A)** Funnel plot for the assessment of publication bias for the effects included in the quantitative analysis. Effects of each included study are presented by dots. The size of the observed effects (Hedge’s *g*) is mapped on the x-axis, whilst the y-axis represents the standard error of the reported effects. The displayed distribution as well as the significant rank correlation test (Kendall’s **τ** = 0.47, *p* < .01) indicate funnel plot asymmetry, suggesting the presence of publication bias. **B)** Distribution of p-values of all effect sizes included in the quantitative analysis, revealing that the majority of published studies report effects passing the significance threshold – or that studies with significant effects are more likely to be published.

#### 3.2.5 Robustness Control Analyses

To test the robustness of our results against alternative *r-*values used in the correction for dependent measurements (see 2.5 Effect Size Calculation), all analyses were repeated with two alternative *r*-values for the expected correlation between the dependent measures, i.e., *r =* 0.74 (maximal liberal estimate) and *r* = 0.97 (maximal conservative estimate). Both control analyses did not change the results of the global effect model markedly (see Supplementary Material C Fig. S2a and Fig. S3a). Also, all but one subgroup model (the downregulation model) remained significant in the liberal correction mode. For the conservative mode, all models but two (the downregulation model and the model for individual frequencies) remained significant. More detailed results of these control analyses are presented in Supplementary Material C Fig. S2 and Fig S3.

To conclude, the results of these control analyses suggest that the correction used for dependent measures (i.e., the estimated *r-*values for the correlation between two dependent measurements) did not affect the outcome of the outlier and influence analysis in the context of the global model. For the subgroup analyses, however, the *r-*values used for the correction of the dependent measures could have biased which effects were considered as influential outliers (the changes are reported in Supplementary Material C).

## 4. Discussion

With this qualitative review and quantitative meta-analysis, we provide a systematic evaluation of previous research applying EEG-based neurofeedback to modulate frontal midline theta. Further, we present detailed insights into existing methodological differences, potential moderators, and ideas for future developments. Our results propose frontal midline theta neurofeedback as a promising approach to modulate brain activity and human behavior, however, convincing evidence for its effectiveness is still missing. Enormous cross-study heterogeneity, small sample sizes, and different definitions of the term “effective modulation” hinder systematic cross-study comparisons and highlight the pressing need for the development of common standards and larger studies including replication efforts.

### 4.1. Quality of Original Studies

While the decision to stick to our previously defined (see preregistration), but rather strict exclusion criteria, reduced the number of studies eligible for this review and meta-analyses, it assured that the finally included publications were of sufficient quality and showed overall a low risk of bias in study design. This was confirmed by the results of the Study DIAD. The comparably low rating in the category “generalizability of findings” matched expectations, at least for multi-session protocols. To overcome this problem, testing more subsettings and subgroups is essentially required. However, this would also require more participants, which is often a question of funding and feasibility. Additionally, insufficient reporting of outcomes (effect sizes mostly missing) was also identified as critical issue, which needs to be urgently addressed in future research. Notably, our strict exclusion criteria did not decrease the heterogeneity of included study designs highlighting again the lack of standards. With our systematic review and meta-analyses we hope to contribute to the development of such common standards and, thus, to enhance the comparability of different FMT neurofeedback studies and their respective results.

### 4.1. Qualitative Review: FMT Modulation Affects Memory Performance

Our literature review provides across-study evidence for the assumption that successful FMT upregulation is possible within and across neurofeedback sessions, which, to some extent, is confirmed by our subgroup meta-analysis. For downregulation, no clear picture evolved, with only one study (Chen et al., 2022) reporting significant changes in the desired direction.

#### 4.1.1. Responder Rates

Critically, only three of 14 original studies reported responder rates. Even though the reported numbers were in line with neurofeedback studies addressing other neural functions (e.g., alpha in visual brain areas; for review see Alkoby et al., 2018), this is highly problematic and the fact that no common definition of the term “responder” exists, complicates the interpretation further. The percentage of people who will likely be able to modulate the desired neural function directly impacts the feasibility and efficacy of the intervention and is, therefore, important to know in advance, i.e., during study planning, as well as for proper interpretation of the study results. Therefore, we here developed a common nomenclature and recommend its use in future research (see below, section 4.3. Future Direction and Recommendations for Future Research).

#### 4.1.2. Modulation of Behavior

Changes induced on task-related EEG were not meta-analytically summarized, but the six studies included in our narrative review reported heavily varying results. This is not surprising, considering the different tasks employed and the intended modulation of different cognitive processes, some of them may be easier to modulate than others. Thus, as long as the low number of available studies hinders task-specific meta-analytic comparisons, replicating previous studies is extremely important to gain a realistic estimate of the underlying effect.

Most successful behavioral alterations induced by FMT neurofeedback were observed in the domain of memory manipulation, which is in line with a recent meta-analysis suggesting that episodic and working memory can be improved by FMT neurofeedback training (Yeh et al., 2022). Less well evidenced are modulations of inhibitory control, motor learning and changes in sports performance. Overall, the results of our review are complemented by basic research linking frontal theta oscillations to memory functions (Hsieh and Ranganath, 2014; Tóth et al., 2014), and provide a first basis for possible applications in a therapeutic context (e.g., for treating dementia).

#### 4.1.3 Long-term Effects

Concerning the investigation of potential long-term effects, one study observed that some changes in behavior and event-related desynchronization were strongest 13 days after the last neurofeedback session (Eschmann and Mecklinger, 2022). The observation of delayed effects is not unusual. Similar delays (across-study effects were most pronounced after 2-12 months) have, for example, been demonstrated in a meta-analysis investigating the impact of neurofeedback on ADHD symptoms (Van Doren et al., 2019) and in fMRI neurofeedback studies (e.g., Rance et al., 2018). These observations imply the possibility that in neurofeedback studies using a more restricted time frame or include no follow-up measurement at all, the strongest effects of an intervention could be missed. Potential reasons for such delayed effects include the integration of learned modulation strategies into daily life and therefore a quasi-unmonitored continued practice as well as physiological changes induced by long-term consolidation processes. Even though both explanations might be valid and even complementary, Gevensleben et al. (2014) provide interesting reasoning for the latter by proposing that neurofeedback leads to unconscious changes in a so-called “EEG-trait” (a sustained change in the oscillatory brain activity), while the use of strategies would imply the “acquisition of a skill”, which would rather more be linked to momentary changes in the “EEG-state”.

In the light of long-term effects, it is notable that the lengths of the here reviewed interventions were fairly short as compared to successful clinical interventions, which would usually incorporate 20-40 sessions per participant (Hammond, 2011) as well as in comparison to the beginning of neurofeedback where participants trained sometimes for more than 100 hours (Birbaumer et al., 1999; Kübler et al., 1999; Lantz and Sterman, 1988; Sterman, 1977). The here included studies did not exceed 12 sessions per participant, with four of them trained only for a single session. While the optimal number of sessions remains a matter of debate, it surely will affect the outcome of the intervention. However, the current research landscape, shaped by funding policies and the pressure to rapidly publish, poses challenges to a comprehensive examination of this question. Hence, similar to numerous other scientific fields, neurofeedback research calls for the adoption of “Slow Science”, advocating for more deliberate and considered research practices (Gärtner et al., 2024; Stengers, 2016).

### 4.2. Quantitative Meta-Analyses Suggest Medium Effect Sizes and Identifies Possible Moderators

Our random effects model for the global effect of FMT neurofeedback revealed a significant across-study effect of medium size. Despite this effect was reduced to trend-level after the influential outlier was removed, it provides at least preliminary evidence for the trainability of FMT modulation. A more robust across-study effect was found when only the modulation runs from the beginning to the end of a session/intervention were compared (contrast 2). This suggests increased changes of modulation across time, most likely caused by learning. Further, our meta-analytical findings provide empirical evidence for multi-session protocols being more successful than single session protocols as well as for the circumstance that downregulation of FMT is harder to achieve than FMT upregulation. However, results of the downregulation model, although significant, should be interpreted with caution, as across-study heterogeneity in the reported effects was very high and most of the included effects stem from a single study with a single session protocol (Chen et al., 2022).

#### 4.2.1. Specific Considerations for FMT Inhibition

When globally considering all original study effects, it is striking that all four observations of negative effect sizes (modulation effects against the intended direction) stem from studies with single session protocols aiming at FMT downregulation (Chen et al., 2022.1, Chen et al., 2022.2, Rozengurt et al., 2017.1, Shtoots et al., 2021.2). Two reasons might be considered here.

##### 4.2.1.1. Number of Sessions

First, a single training session might simply be not sufficient to induce the intended effects. This assumption is supported by the observation that most and on average stronger effects in the intended direction were reported by studies employing multi-session protocols.

##### 4.2.1.2. The Modulation Dilemma of FMT Inhibition

Second, FMT downregulation might in general be more difficult to entrain than FMT upregulation. Increased FMT has been associated with increased self-regulation needs (Emmert et al., 2016; Ninaus et al., 2013), conflict detection (Cavanagh and Shackman, 2015; Nigbur et al., 2011), reward processing (Amiez et al., 2012) and, in general, with internally directed cognition (Cavanagh and Shackman, 2015). Such processes might also be induced by the experimental setting itself. For instance, self-regulation is required for the modulation of a brain state, a conflict may arise at the beginning of the neurofeedback training when participants cannot yet relate the feedback they receive to a specific mental strategy, and positive feedback can be perceived as a reward. These processes can unintentionally interfere with the learning process in both directions. In protocols striving for FMT upregulation, the intended and to be positively reinforced brain state (increased theta) may be present more often than without such interfering influences, while in FMT downregulation protocols, the intended brain state (decreased theta) may be present more seldom – similarly the respective feedback signal. While the former would facilitate the learning process, the processes inherent to FMT cause the following dilemma for the latter: Whenever a participant is asked to downregulate FMT and invests efforts to do so, this induces a self-regulation need and facilitates conflict detection, i.e., two processes that by themselves increase rather than decrease FMT. This ultimately induces contradictory influences and induces conflict.

Thus, due to the involvement of FMT in executive functioning and cognitive control, FMT neurofeedback might represent a type of neurofeedback for which it is especially difficult to draw conclusions about the specific underlying processes. Implicit learning could be one solution to overcome the involvement of cognitive control mechanisms, e.g., protocols in which participants are not explicitly instructed to modulate their brain activity and feedback is only provided in disguise so that cognitive load and self-regulatory demands are reduced (Muñoz-Moldes and Cleeremans, 2020). First evidence for the feasibility of “passive covert feedback” protocols was recently provided by a fMRI-neurofeedback study aiming at modulating functional connectivity (Ramot et al., 2016). Nonetheless, this does not solve the issue of involved reward processing, which is necessary even for the simplest forms of operant conditioning (i.e., implicit learning), but also plays a role when learning a mental strategy (i.e., explicit learning).

#### 4.2.2. Across Study Effect Diminished by Study Heterogeneity

Regarding the meta-analytic synthesis in general, it is not surprising that no strong across-study effect was observed given the variety of study designs, the small sample sizes, and the diversity of the reported effects defined with heterogeneous nomenclatures. This high heterogeneity among studies also complicates the interpretation of the publication bias, as the observed asymmetry could also result from different study design choices. Two of such differences in study design are exemplary discussed below:

##### 4.2.2.1 Multiple Frequency Bands

For one, those studies whose feedback signal reflects a ratio of FMT and another frequency band might provide participants with a shortcut to positive feedback. They receive positive feedback not only when FMT is modulated in the intended direction (e.g., increase), but also when it moves in the opposite direction (e.g., decrease), as long as the other frequency band changes are stronger than the changes in FMT. This may be comparable to trying to reduce caffein intake by reinforcing thinly brewed coffee. It is quite easy to imagine how one could just start drinking more coffee, i.e., increasing the amount of water instead of decreasing the amount of coffee powder and therefore the overall consumed caffeine is similar. If it is easier for the participant to control the other frequency, the necessity to control theta is undermined and increases the probability for null- or negative effects.

Even though it has been proposed that targeting more than one band during neurofeedback may be difficult to entrain (Rogala et al., 2016), feedbacking multiple frequency bands or ratios can be advantageous under several circumstances. For example, one of the best evidenced neurofeedback therapies is the one for ADHD, which commonly targets the theta/beta ratio. Depending on the goal of the neurofeedback intervention, a ratio may help to reduce band interdependence or to achieve a specific brain state consisting of multiple frequency components.

##### 4.2.2.2 Control Conditions

Secondly, effects across subjects highly depend on the applied control conditions, which also differed among studies. Sham-neurofeedback was the most frequently used control condition in the meta-analytically compared investigations. However, this might not be an optimal choice, especially in FMT neurofeedback, as self-regulatory processes and conflicts induced by the experimental situation itself may affect FMT but not (or differently) the other frequency band, ultimately leading to the observation of a spurious effect (Davelaar et al., 2018). But as already discussed in the literature (Ros et al., 2020; Sorger et al., 2019), different control conditions are characterized by different problems.

#### 4.2.2. Publication Bias

All results of our publication bias analyses support the presence of serious publication bias in the field of FMT neurofeedback. Specifically, our results suggest that studies with larger standard errors, which is typically the case in studies with small samples, are more likely to report larger effect sizes – and to ultimately get published. Although, it is crucial to note that these indicators of publication bias are limited by the small number of studies available (especially effects investigating FMT downregulation as well as effects across subjects are underrepresented in our sample), we must be aware of the fact that the here reported across-study effect size most likely represents an overestimation of the true effect. In the future, this problem can be overcome by the strict implementation of Open Science practices in the field of FMT neurofeedback such as the preregistration of hypotheses and analyses plans or the use of registered reports that facilitate the publication of null- or contradictory findings and do thus contribute to a more realistic picture of the true effect (Lotte et al., 2019).

In sum, even though small sample sizes, diverse study designs, and highly heterogeneous effects render a review and even more a meta-analytic synthesis of across-study findings on FMT neurofeedback challenging, we here reveal that FMT modulation is possible. However, whether a FMT neurofeedback intervention is successful may critically depend on choices in study design characteristics, which need to be thoroughly explored and validated (see next paragraph).

### 4.3. Future Direction and Recommendations for Future Research

The problems and challenges which arose during our synthesis corroborate previous critiques of neurofeedback studies. However, given the high costs and efforts associated with neurofeedback studies, it will continue to be difficult for researchers to address them with an adequate sample size and study design, e.g., proper control conditions and enough sessions, with the current research policies in place.

Nevertheless, as reliable insights into underlying processes and into the effectiveness of interventions may only be derived from multiple studies pointing into the same direction, and especially against the background of the current “replication crisis” in psychology (Wiggins and Christopherson, 2021), it is today even more important than ever to reach transparent and reproducible research practices (Arslan, 2019; Polanin et al., 2020). One important way towards this goal is standardization, based on insights from meta-analyses and systematic reviews (Mueller et al., 2018). Although efforts and costs may be non-negligible, we believe the field can only move forward, if we thoroughly investigate different design choices, follow proposed guidelines, gather evidence and finally summarize the findings in systematic meta-analyses, which provide the basis for the definition of new standards (Ros et al., 2020). We here aim at contributing to this endeavor by disentangling the vast term of “effective modulation” and the following four recommendation to future research:

#### 4.3.1. Defining Responders

The heterogeneity of reported effects, in terms of what is considered as effective modulation, is a problem not easily solved. Critically, even the CRED-nf criteria (Ros et al., 2020), which were explicitly developed to increase study quality and reporting praxis, were not able to provide a clear recommendation in many cases, e.g., concerning the question of with which timepoint (e.g., resting state, first modulation run, etc.) the modulation run(s) should be compared. This is in so far reasonable, as depending on the underlying research question, different effects might be of primary interest. However, a clear and common definition of the term “effective modulation” and a thorough distinction between different kinds of modulation effects are essential in any case. We here propose a possible standard for classifying neurofeedback studies and suggest a common nomenclature for its reporting (Table 2):

**Table 2.** Proposed classification and nomenclature for neurofeedback responses.

| Domain | Type | Subtype | Changes | Description<br>(Timepoints and conditions of compared EEG recordings) | Research Question |
| --- | --- | --- | --- | --- | --- |
| *Feedback* | W |  | State | Changes from the resting-state to the modulation blocks with feedback within sessions | Testing for presence or absence of the modulation |
|  | A | Mod | State | Changes of modulation blocks with feedback across sessions | Testing for training effects of modulation |
|  |  | Diff | State | Changes in the difference between the resting-state and the modulation blocks with feedback across sessions | Testing for training effects of modulation, accounting for baseline changes |
| *Transfer* | W |  | State | Changes from the resting-state to the modulation blocks without feedback within session | Testing for the effectiveness of mental strategy / voluntary modulation |
|  | A | Mod | State | Changes of modulation blocks without feedback across session | Testing for training effects of mental strategy / voluntary modulation |
|  |  | Diff | State | Difference between the resting-state and the modulation blocks without feedback across session | Testing for training effects of mental strategy / voluntary modulation, accounting for baseline changes |
| *Resting State / Task* | I |  | State | Changes in resting-state or task immediately after a neurofeedback session | Testing for short-term effects after neurofeedback |
|  | C |  | Trait | Changes in resting-state or task measured across multiple sessions, measured before the modulation (e.g., baseline recordings) | Testing for long-term effects after neurofeedback |
|  | F |  | Trait | Changes in resting-state or task measured at least several hours after the last neurofeedback session | Testing for long-term effects after neurofeedback |

Note: Recommendation for a uniform classification and naming of different modulation effects to increase the comparability of neurofeedback studies. A = across session, C = continuous, D = difference, F= follow-up, I = immediate, Mod = modulation block, W = within session.

##### 4.3.1.1. Domains

We suggest the distinction of three kinds of responders (*domains*):

1. “feedback responders” show altered brain activity during modulation runs in which feedback is provided,
2. “transfer responders” demonstrate altered brain activity during modulation runs in which no feedback is provided, and
3. “resting-state / task responders” show altered brain activity during resting state as compared to during task performance compared to measurements before the intervention.

##### 4.3.1.1 Types

Further, we define five *types (I-V)* of responders depending on the scope of the observed changes:

- For “feedback and transfer responders” (domain 1 and 2) we suggest:

(I) the “type W” describes a change within a session, and

(II) the “type A” refers to a change across sessions.

The “type A” may be further subdivided in two *subtypes*:

(IIa) “type A_mod” for comparisons between modulation runs across different sessions, and

(IIb) “type A_diff” for comparisons between the resting state-to-modulation difference across sessions.

- For “resting-state / task responders” (domain 3) three further types were defined:

(III) “type I” defines changes immediately after a neurofeedback-session,

(IV) “type C” for changes observed continuously across several neurofeedback sessions or during the resting state before the modulation, and

(V) “type F” for changes observed in follow-up sessions.

Of note, “Type A_mod” will not account for resting-state changes, as measured with “type C”, “type A_diff” will do so. Further, while changes immediately after neurofeedback-sessions (III - “type I”) may still depend on induced plasticity and might therefore present a state change, the latter changes (IV-“type C” and V-“type F”) are likely to represents long-term alterations, i.e., a trait.

#### 4.3.2 Data Sharing

Comparing effects across different studies represents only one means to gain a more realistic estimate of the effectiveness of FMT neurofeedback. However, the above demonstrated diversity of nomenclatures, study designs, and especially the lack of proper reporting of effect statistics hinder systematic across-study comparison. Therefore, we strongly advise future studies to provide at least means and standard deviations for all conducted measurements, either in the main text or in the supplementary material. A complementary means to gain more realistic estimates is to increase statistical power by enlarging sample sizes. Given the high efforts and cost for neurofeedback-studies, we consider it therefore as indispensable for neurofeedback researchers to collaborate, to share their code and data, and to build up multi-center projects.

#### 4.3.3 Preregistration of Planned Analysis

Another issue posed by the high variety of effects is the danger of selective reporting and “cherry picking”, i.e., only reporting results that best fit to the posed hypotheses (Andrade, 2021; Chambers et al., 2014). Consequently, we deem it mandatory to preregister not only study designs but also all details (e.g., the statistical test, its parameters, *p*-thresholds, etc.) of the intended preprocessing and analysis pipeline. Importantly, this should not preclude authors from conduction exploratory analysis but enforces that those will be marked as such.

#### 4.3.4 Multiple Sessions and Follow-Ups

Further, we would like to advise researchers to include multiple sessions in their protocol, especially those who are aiming at FMT inhibition, and to implement a follow-up assessment. Training participants for much longer time periods, similar to early days of neurofeedback, may allow for (stronger) manifestations of potential effects (Birbaumer et al., 1999; Kübler et al., 1999; Lantz and Sterman, 1988; Neumann et al., 2003; Sterman, 1977). Further, learning processes can also take place after finishing the last session of an intervention. These would be only detectable in a follow-up assessment taking place after weeks or months. To assess both brain activity and potential behavioral changes across a longer time frame, would allow for important insights into the learning processes of FMT neurofeedback that might be missed otherwise.

#### 4.3.5 Insights from Other Domains of Research

While this meta-analysis primarily focuses on continuous frequency band training in EEG-neurofeedback studies, Batail et al. (2019) stress the significance of basing neurofeedback on specific well-known biomarkers or on solid neurophysiological principles (e.g., EEG coherence, Li et al., 2019; event-related brain potentials (ERPs), Cavanagh and Frank, 2014; Cavanagh and Shackman, 2015). However, in contrast to protocols with continuous feedback, ERP-based protocols would require intermittent feedback on the basis of specific trials, which overall implies that less frequent feedback is provided. Nevertheless, a learning curve for increasing ERP amplitudes after training has been demonstrated in the field of brain-computer interfacing (BCI, Eidel and Kübler, 2022; Ziebell et al., 2020) and, in general, observations from BCI research may have great potential to be transferred to the domain of FMT-neurofeedback, as already suggested by Jeunet et al. (2018). Specifically, they propose the use of adaptive thresholds, machine learning tools for customizing neurofeedback features, applying performance predictors for selecting suitable neurofeedback protocols, integrating tactile and computerized social and emotional feedback to boost motivation, and the identification of optimal mental strategies through algorithms initially developed for BCI to enhance individual training outcomes.

To sum up, for neurofeedback in general we recommend the use of the here proposed nomenclature to clearly declare which domain(s) of modulation is/are intended, which one(s) is/are the one(s) reported, and to provide the respective responder rates. Further, we encourage consideration of the CRED-nf (Ros et al., 2020), to openly share data with other researchers, to conduct follow-up assessments and to incorporate insights from other domains of (neurofeedback) research. For frontal midline theta neurofeedback specifically, we advocate the application of multiple sessions and to more thorough investigation frontal midline theta downregulation.

### 4.4. Limitations

The first major drawback of our meta-analysis is the availability of only few original studies that fulfill the inclusion criteria and, thus, could be included in the cross-study comparison. This circumstance forced us to extract data from whatever source available (see section 2.4. Effect Size Extraction), such that the second limitation refers to potential distortions of effect sizes which may have occurred during the conversions, calculations, and extraction of effect sizes from graphs (see section 2.5. Effect Size Calculations). However, given the lack of reported means and standard deviations, this was the only way possible to address our research question.

Third, the small number of included studies prevented also the inspection of further variations in study design details. Enriquez-Geppert et al. (2017) describe in detail the different parameters which need to be set when designing a neurofeedback study and optimal choices for some of these parameters have been assessed by different studies, but do not always align. For example, a direct comparison of auditory against visual feedback suggested favoring visual feedback for the modulation of slow cortical potentials (Hinterberger et al., 2004). On the other hand, summaries of other reports, could not confirm the superiority of one modality over the other, but rather suggest a combination of both (Rogala et al., 2016; Vernon et al., 2009, 2004). Despite these efforts, reliable optimal settings or choices for these parameters are not yet well evidenced (Rogala et al., 2016). This renders each of these parameters an interesting target for subanalyses and further investigation. Here, we decided to focus on those parameters for which we expected (at the point of our preregistration) the strongest influence on modulation and for which we assumed to have enough valid data. Additional interesting analysis when sufficient studies are available, would, for example, include the feedback modality (see Rogala et al., 2016), the session duration and number or the type of instruction and strategies communicated to the participants (see Kober et al., 2013 or Sepulveda et al., 2016). Even though we could not systematically address this issue in our meta-analysis due to the few available studies, Chen et al. (2022) also provided first evidence that function-specific instructions may be necessary for FMT downregulation.

The fourth difficulty introduced by the heterogeneous study designs is the possibility that some observed effects reflect different underlying true effects which are of opposite directions. If such effects are combined within the same model, the model will likely predict a null effect, even though two discernible ones are present. This is especially problematic as the underlying mechanisms of neurofeedback learning are far from being fully understood. Most critical, it is often unclear if adaptions in resting-state and during task-related EEG occur in the same (i.e., Hebbian plasticity) or opposite (i.e., homeostatic plasticity) direction as in the direction trained during the neurofeedback. If some parameter settings would induce the first kind while others induce the latter, it would increase the possibility that effects may level out.

Overall, the limited amount of available data has significantly influenced the level of granularity at which we could investigate across-study effects. Only future meta-analyses based on studies higher in number and with full reporting of all relevant data can remedy this issue.

## 5. Conclusion

We conducted a systematic review and quantitative meta-analysis to summarize the current empirical evidence on the efficacy of frontal midline theta neurofeedback. Results of 14 studies were summarized in a systematic review, while 17 effects extracted from 11 studies were synthesized in a quantitative meta-analysis. Our review demonstrates that the most notable behavioural improvements following FMT neurofeedback are linked to memory enhancement, while effects on inhibitory control, motor learning, and sports performance remain less substantiated. Our quantitative meta-analysis revealed a significant across-study effect of medium size for frontal midline theta modulation via neurofeedback in general. Across-study effect sizes for subgroup analyses were also of medium to even large size, but nearly all meta-analytical comparisons demonstrated significant across-study heterogeneity in the reported effect sizes. The results of the conducted subanalyses suggest that a) changes are most prominent during modulation runs from the beginning to the end of a session/intervention, and b) that single session protocols, especially for FMT inhibition might be less effective. Even though the heterogeneity of the available data did not allow drawing strong conclusions, we provide an overview of the current state of the art, which may serve as basis for informed decisions in future research. Finally, we propose a common nomenclature and classification system for neurofeedback research that may facilitate future across-study comparison.

To conclude, more research is urgently required to validly reveal how FMT may be modulated by means of neurofeedback. This research needs to be preregistered, reported in an adequate manner, and data to be shared to enable a proper synthesis of results. Assuming small sample sizes as well as diverse study designs will remain a core problem of neurofeedback, a revision of this meta-analysis in a couple of years is essentially required to evidence the capabilities and limitations of FMT neurofeedback and to further refine guidelines for choices in study designs. Funding of long-term studies and acknowledging such efforts in the careers of researchers conducting these studies could contribute to improve future studies and to foster scientific insight into the effects of not only FMT-related but neurofeedback in general.

## Declaration of Competing Interest

Declaration of interest: none.

### CRediT authorship contribution statement

**Maria Pfeiffer:** Conceptualization, Methodology, Formal analysis, Writing – Original Draft, Visualization. **Andrea Kübler:** Conceptualization, Writing – Review and Editing, Supervision, Funding acquisition. **Kirsten Hilger:** Conceptualization, Methodology, Writing – Original Draft, Project Administration, Supervision.

## Acknowledgements

This work was supported by the German Research Foundation [grant number RTG 2660 – Approach Avoidance] assigned to Andrea Kübler. This publication was supported by the Open Access Publication Fund of the University of Würzburg.

## Supplementary Material

Supplementary Material associated with this article can be found in a separate document.

