## Supplement C for "Modulation of Human Frontal Midline Theta by Neurofeedback: A Systematic Review and Quantitative Meta-Analysis"

**SUPPLEMENTARY MATERIAL C**

* Corresponding author:

Kirsten Hilger

Department of Psychology I

Marcusstr. 9-11

D-97070 Würzburg

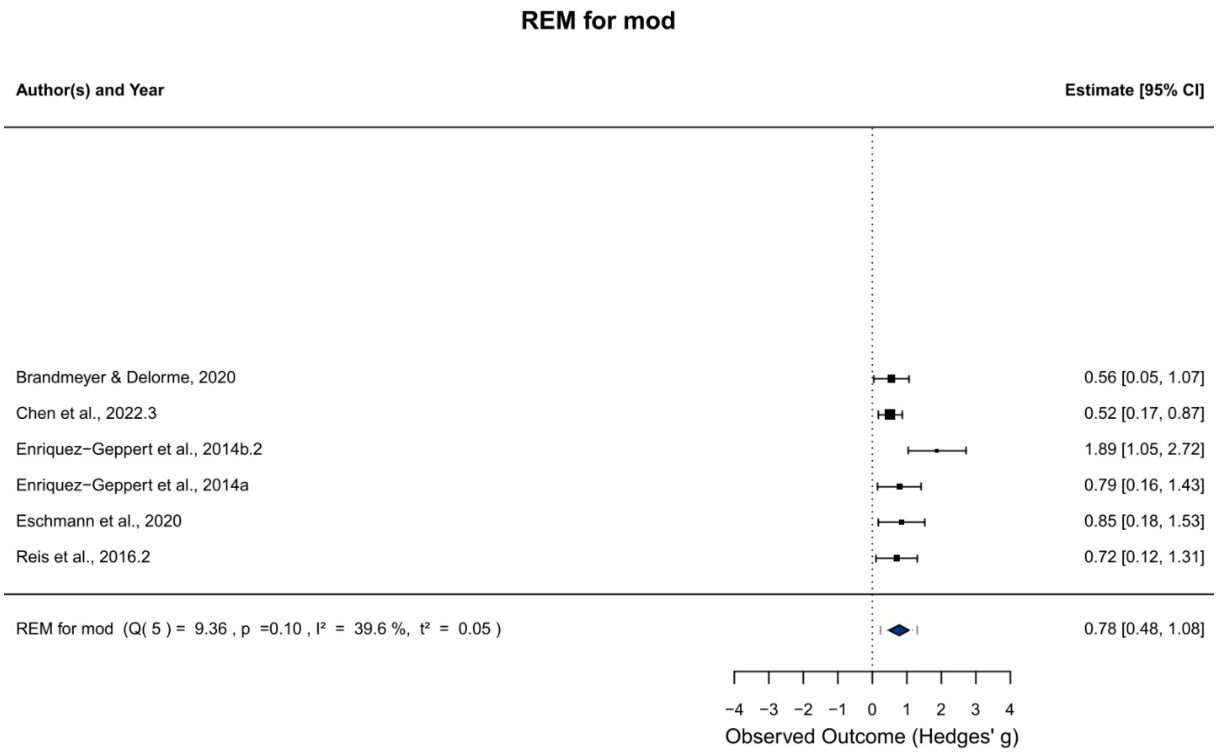


**Fig. S1a** Forest plot for the random-effects model (REM) assessing effects from subgroup (1): “Beginning vs. ending of neurofeedback” – defined as the contrast between either the first and the last block or the first and the last session of an intervention. The black squares denote the mean observed effect sizes, where the size of the square is proportional to the model weight. The 95%-Confidence Interval (CI) is indicated by the whiskers. Diamonds depict the 95%-CI of the REM and their respective prediction intervals (dotted whiskers).

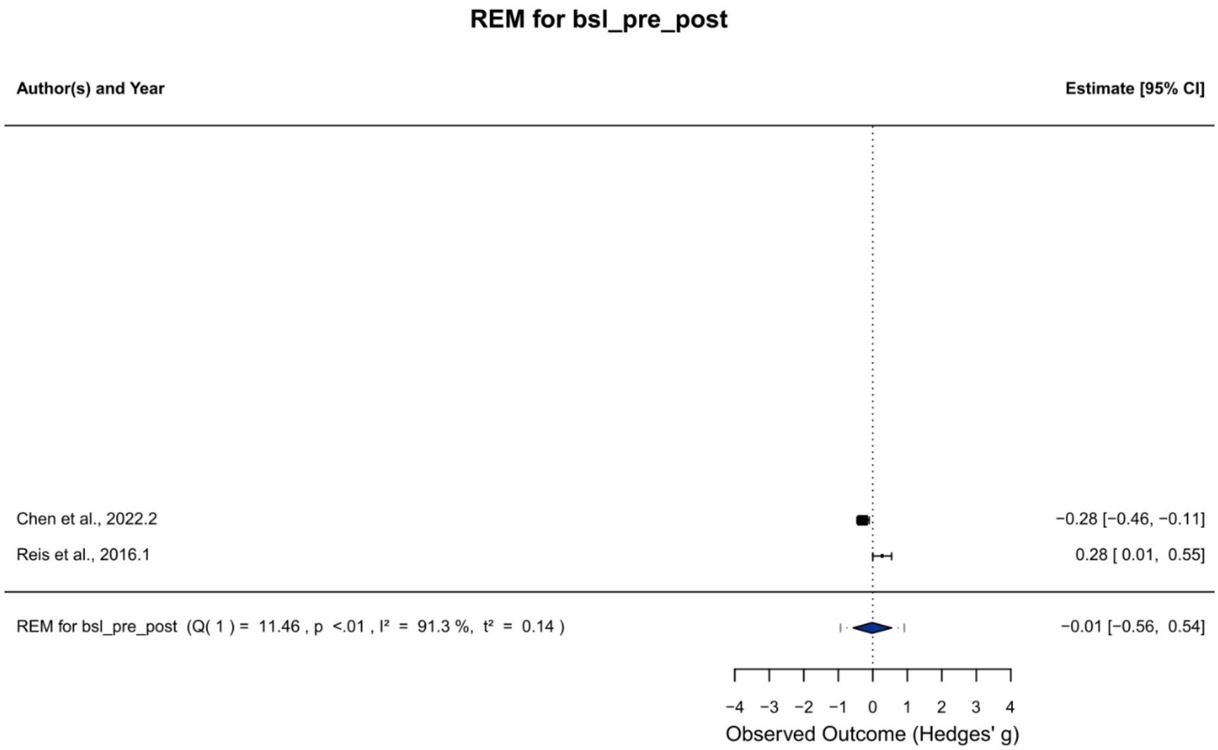


**Fig. S1b** Forest plot for the random-effects model (REM) assessing effects from subgroup (2): “Resting state baseline vs. after neurofeedback” – comparing resting-state measurements before the session to resting-state measurements after a session or to the entire invention. The black squares denote the mean observed effect sizes, where the size of the square is proportional to the model weight. The confidence interval (95%) is indicated by the whiskers. Diamonds depict the 95%-CI of the REM and their respective prediction intervals (dotted whiskers).

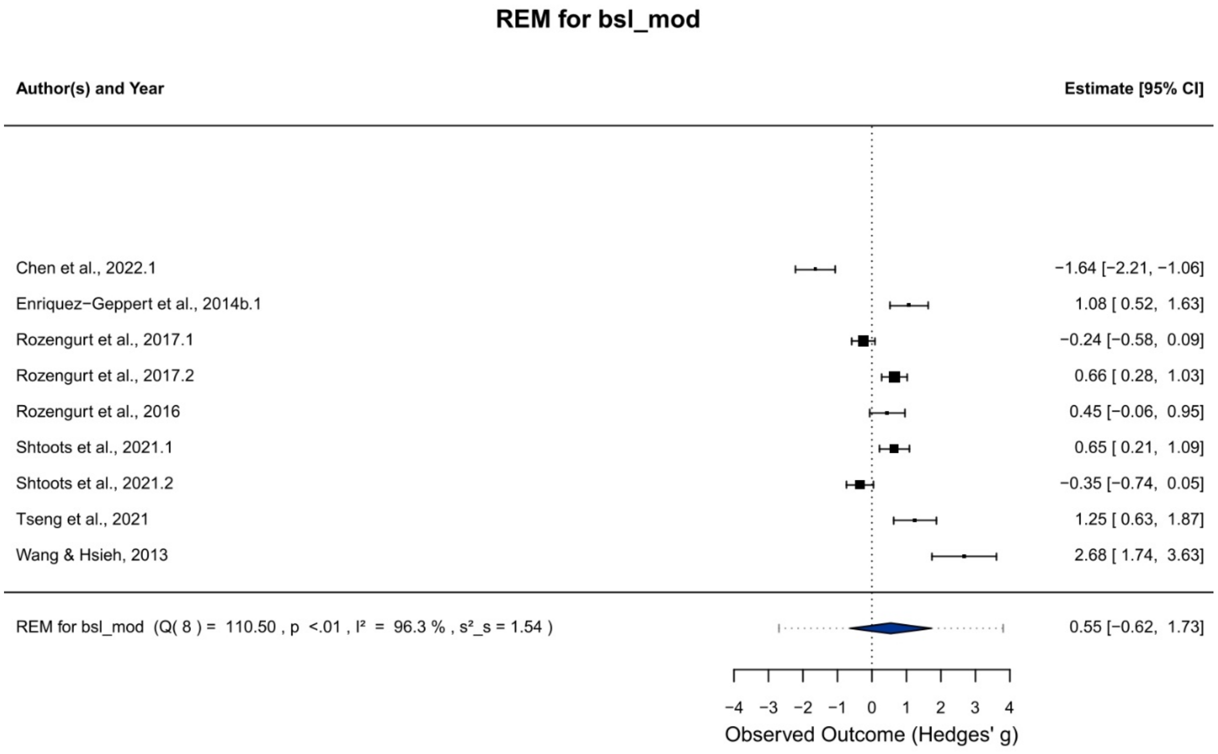


**Fig. S1c** Forest plot for the random-effects model (REM) assessing effects from subgroup (3): “Resting State vs. neurofeedback” – assessing the difference between resting state and blocks in which participants received feedback. The black squares denote the mean observed effect sizes, where the size of the square is proportional to the model weight. The confidence interval (95%) is indicated by the whiskers. Diamonds depict the 95%-CI of the REM and their respective prediction intervals (dotted whiskers).

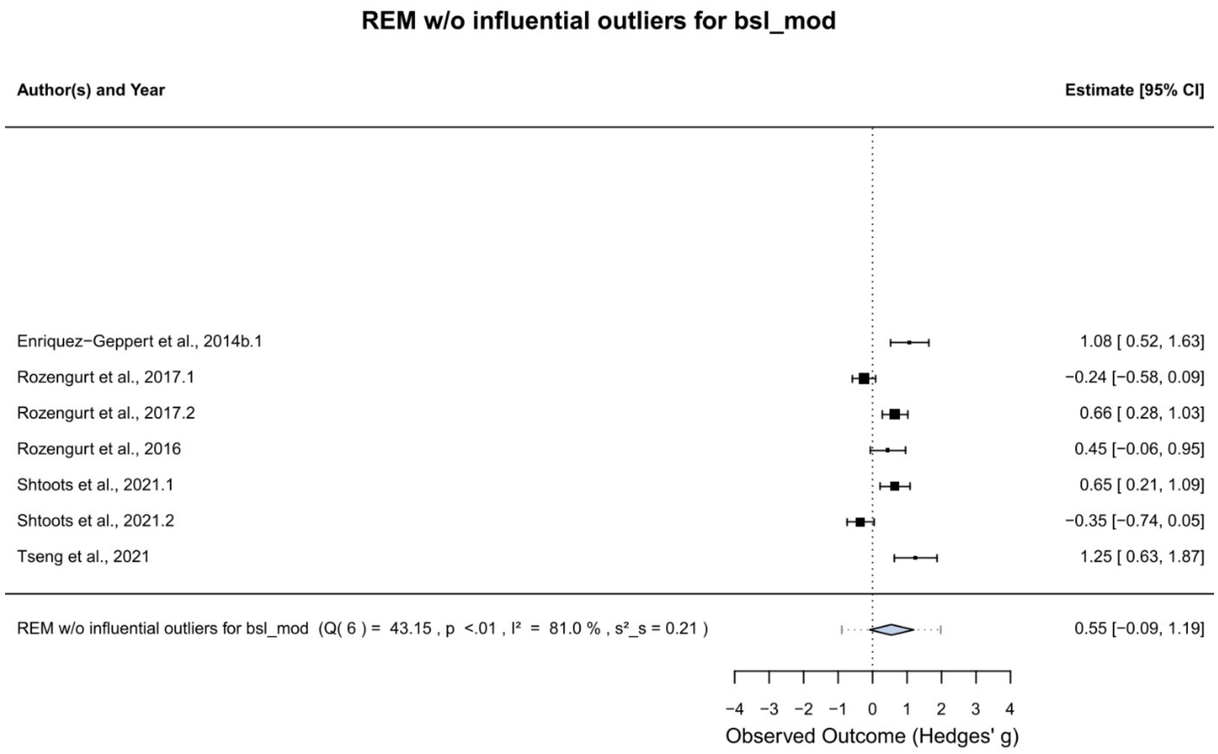


**Fig. S1d** Forest plot for the random-effects model (REM) assessing effects from subgroup (3), without the influential outlier: “Resting State vs neurofeedback” – assessing the difference between resting state and blocks in which participants received feedback. The black squares denote the mean observed effect sizes, where the size of the square is proportional to the model weight. The confidence interval (95%) is indicated by the whiskers. Diamonds depict the 95%-CI of the REM and their respective prediction intervals (dotted whiskers).

**Fig. S1e** Forest plot for the random-effects model (REM) assessing single session protocols.

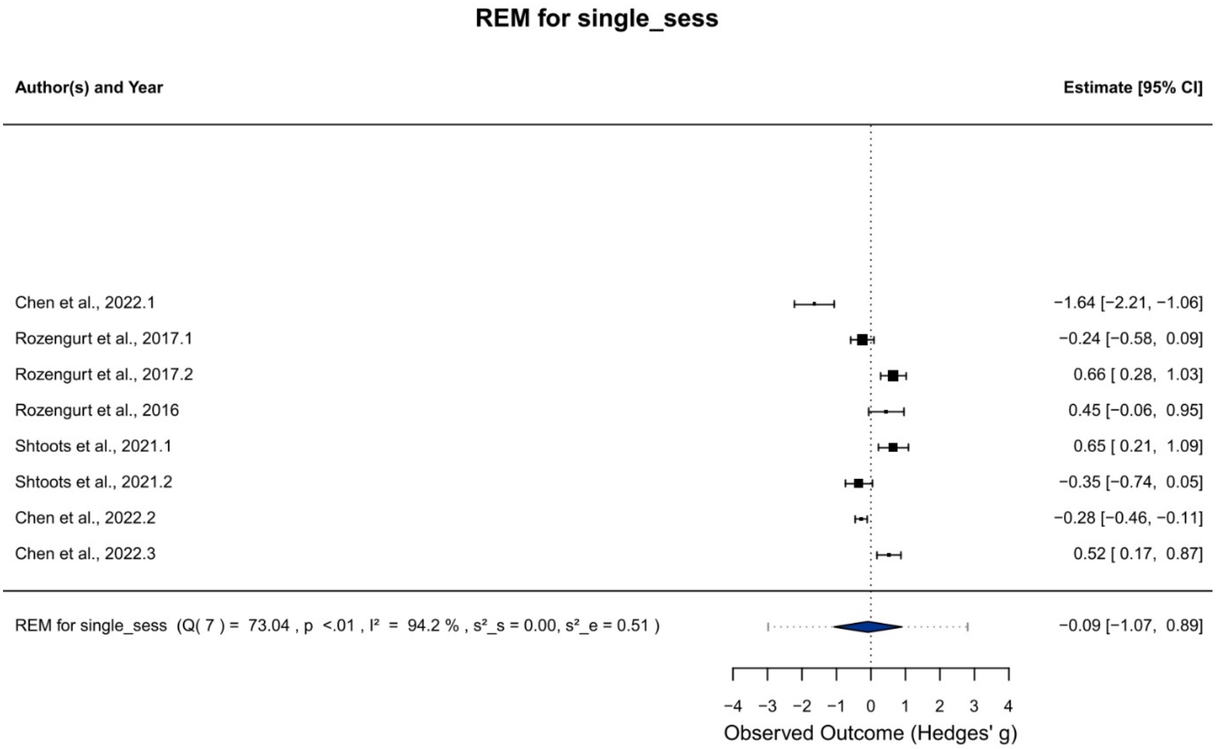


The black squares denote the mean observed effect sizes, where the size of the square is proportional to the model weight. The confidence interval (95%) is indicated by the whiskers. Diamonds depict the 95%-CI of the REM and their respective prediction intervals (dotted whiskers).

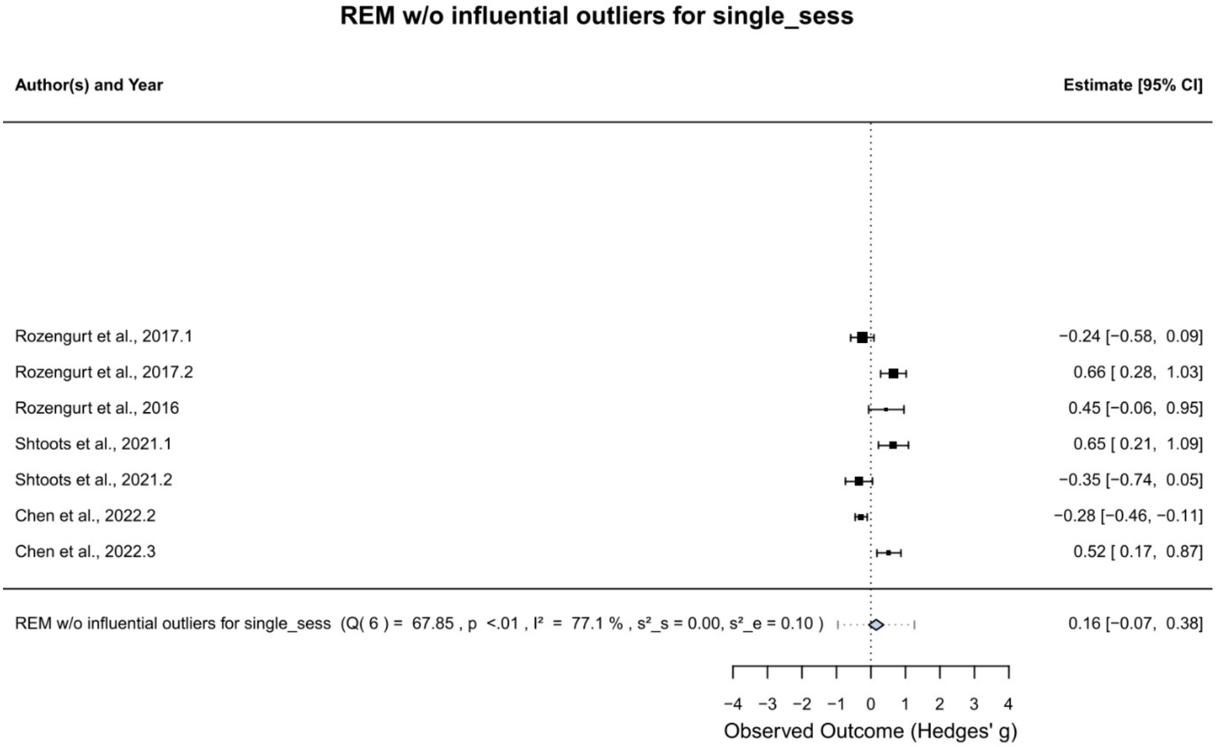


**Fig. S1f** Forest plot for the random-effects model (REM) assessing single session protocols without the influential outlier. The black squares denote the mean observed effect sizes, where the size of the square is proportional to the model weight. The confidence interval (95%) is indicated by the whiskers. Diamonds depict the 95%-CI of the REM and their respective prediction intervals (dotted whiskers).

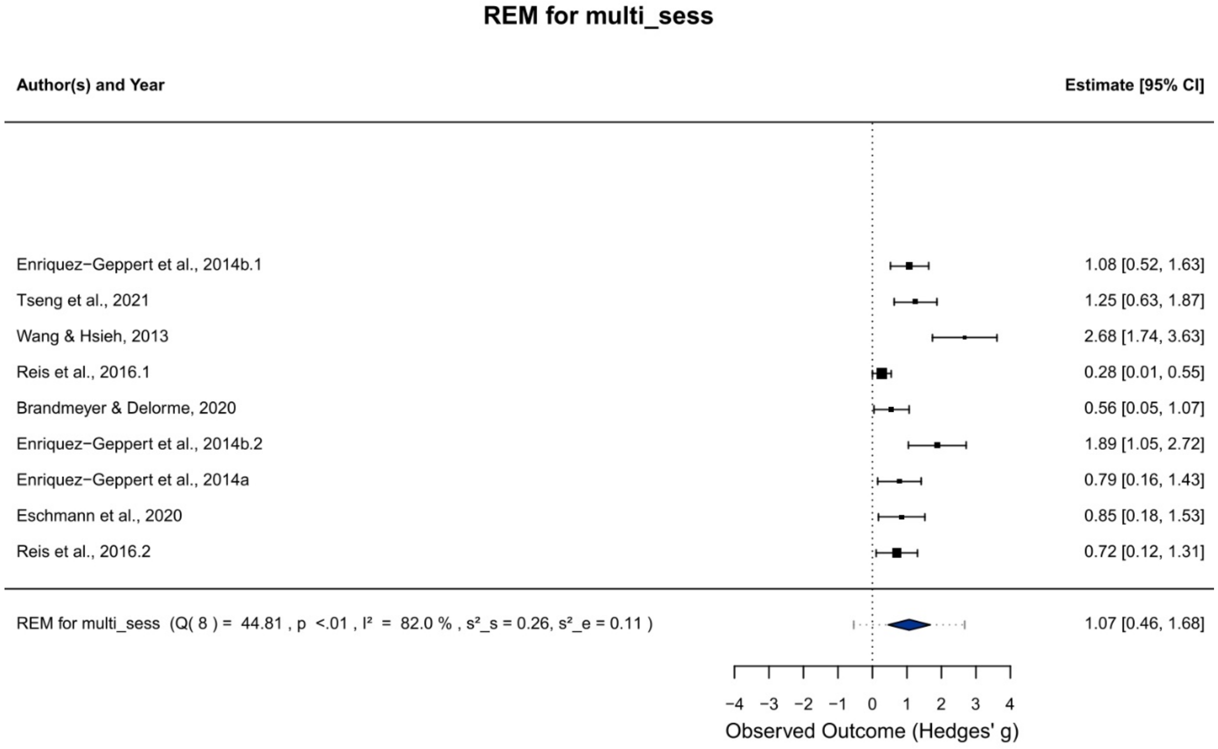


**Fig. S1g** Forest plot for the random-effects model (REM) assessing multi session protocols. The black squares denote the mean observed effect sizes, where the size of the square is proportional to the model weight. The confidence interval (95%) is indicated by the whiskers. Diamonds depict the 95%-CI of the REM and their respective prediction intervals (dotted whiskers).

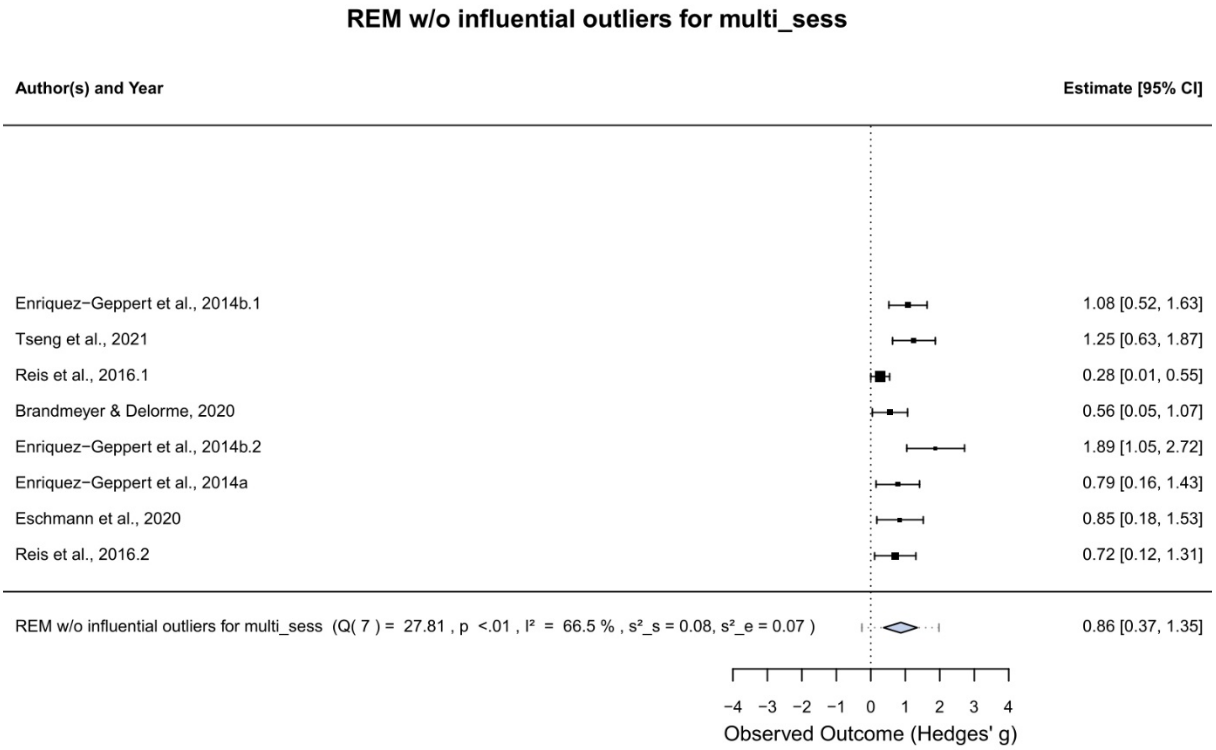


**Fig. S1h** Forest plot for the random-effects model (REM) assessing multi session protocols without the influential outlier. The black squares denote the mean observed effect sizes, where the size of the square is proportional to the model weight. The confidence interval (95%) is indicated by the whiskers. Diamonds depict the 95%-CI of the REM and their respective prediction intervals (dotted whiskers).

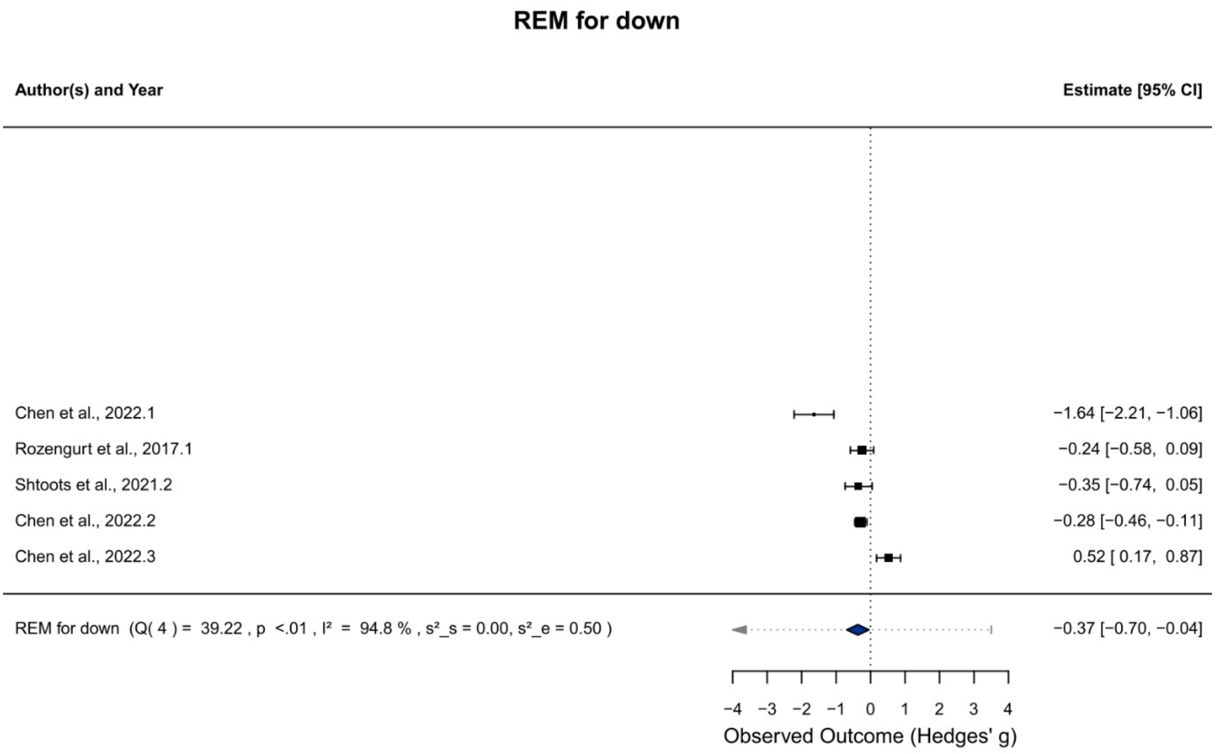


**Fig. S1i** Forest plot for the random-effects model (REM) assessing downregulation protocols. The black squares denote the mean observed effect sizes, where the size of the square is proportional to the model weight. The confidence interval (95%) is indicated by the whiskers. Diamonds depict the 95%-CI of the REM and their respective prediction intervals (dotted whiskers).

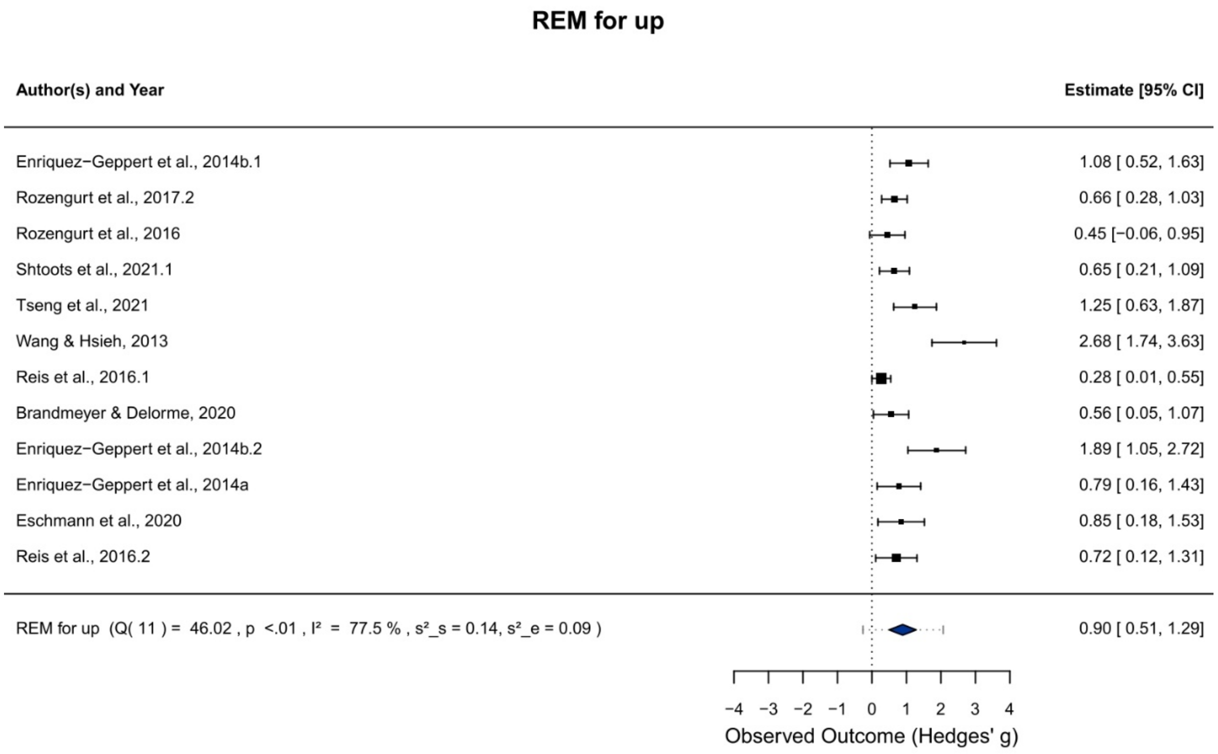


**Fig. S1j** Forest plot for the random-effects model (REM) assessing upregulation protocols. The black squares denote the mean observed effect sizes, where the size of the square is proportional to the model weight. The confidence interval (95%) is indicated by the whiskers. Diamonds depict the 95%-CI of the REM and their respective prediction intervals (dotted whiskers).

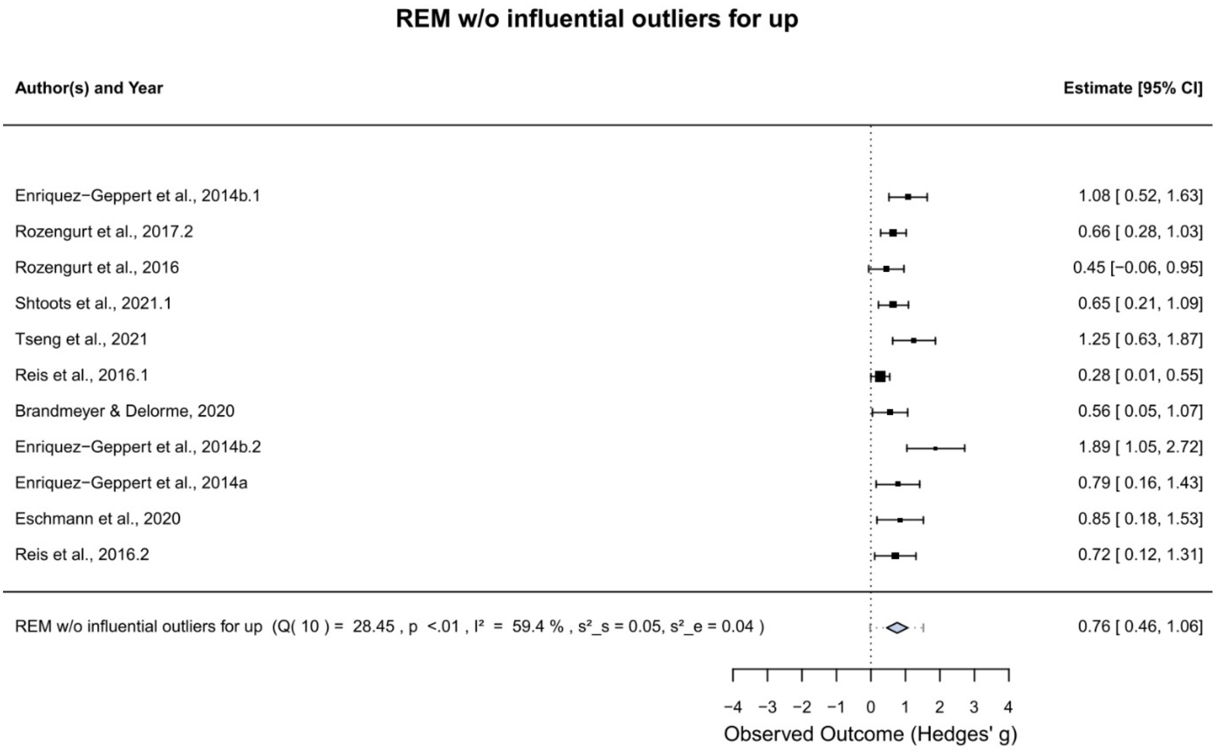


**Fig. S1k** Forest plot for the random-effects model (REM) assessing upregulation protocols without the influential outlier. The black squares denote the mean observed effect sizes, where the size of the square is proportional to the model weight. The confidence interval (95%) is indicated by the whiskers. Diamonds depict the 95%-CI of the REM and their respective prediction intervals (dotted whiskers).

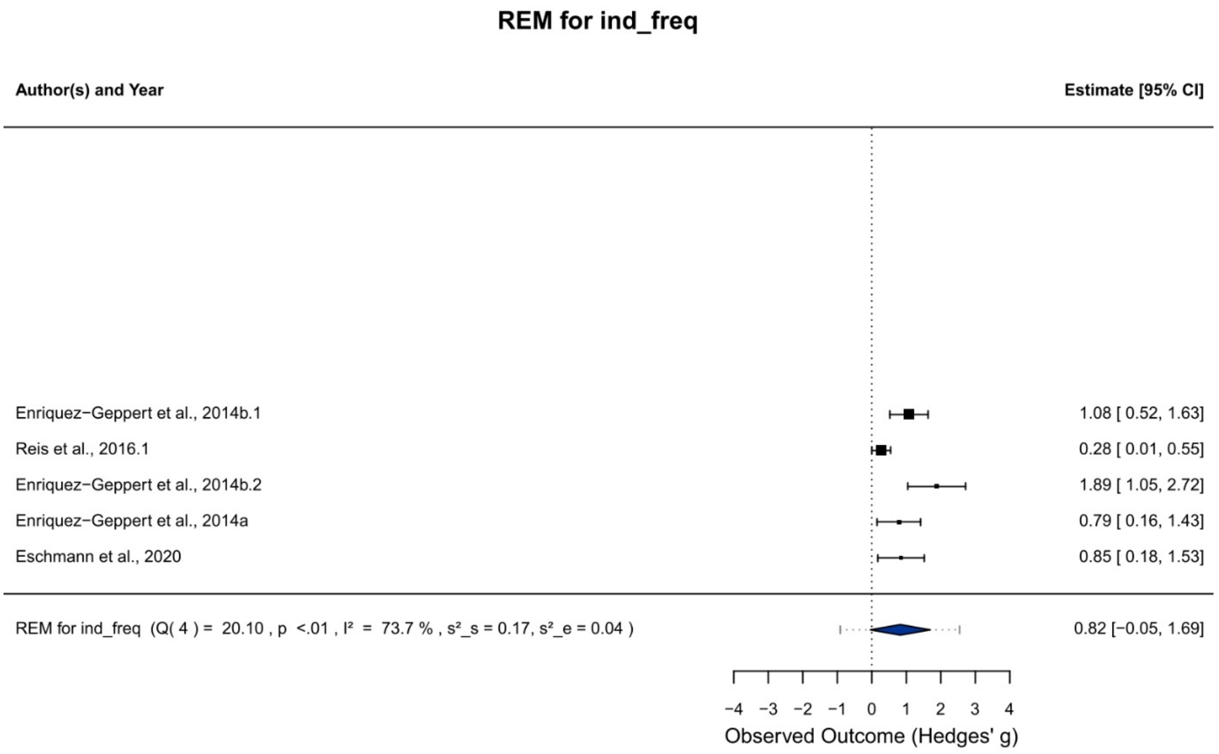


**Fig. S1I** Forest plot for the random-effects model (REM) assessing protocols which used individual frequencies as feedback. The black squares denote the mean observed effect sizes, where the size of the square is proportional to the model weight. The confidence interval (95%) is indicated by the whiskers. Diamonds depict the 95%-CI of the REM and their respective prediction intervals (dotted whiskers).

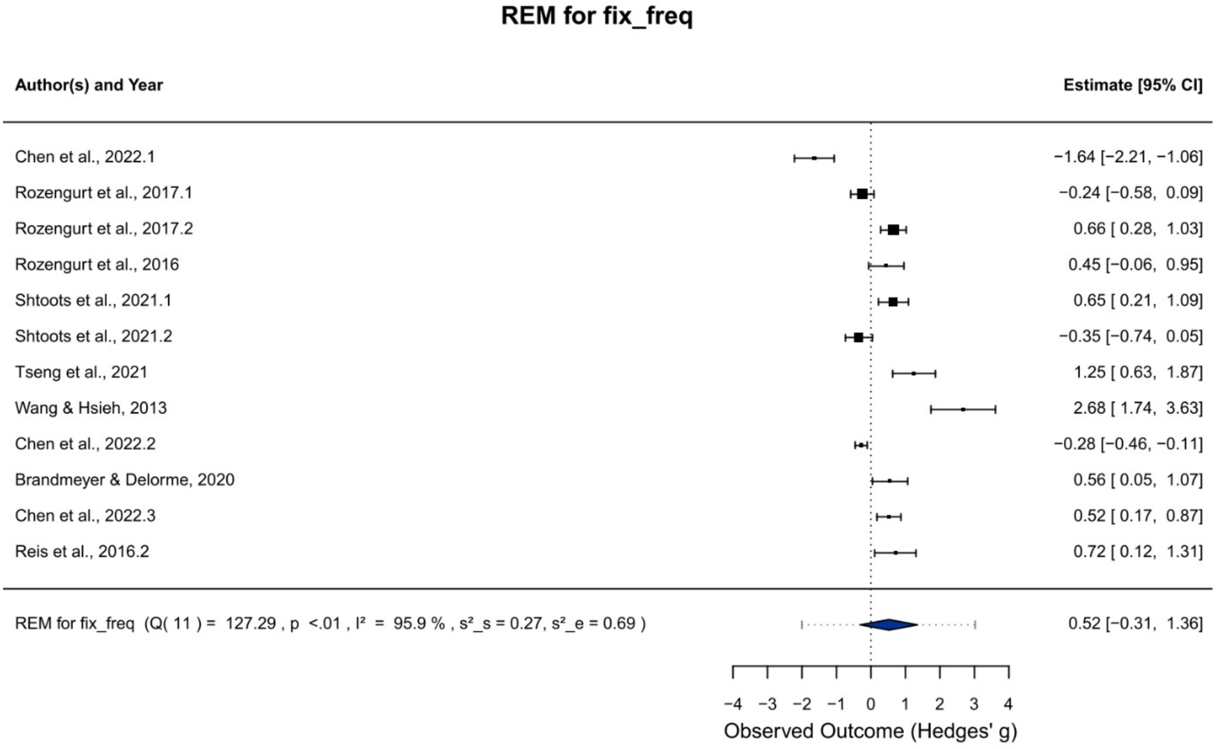


**Fig. S1m** Forest plot for the random-effects model (REM) assessing protocols which used fixed frequencies as feedback. The black squares denote the mean observed effect sizes, where the size of the square is proportional to the model weight. The confidence interval (95%) is indicated by the whiskers. Diamonds depict the 95%-CI of the REM and their respective prediction intervals (dotted whiskers).

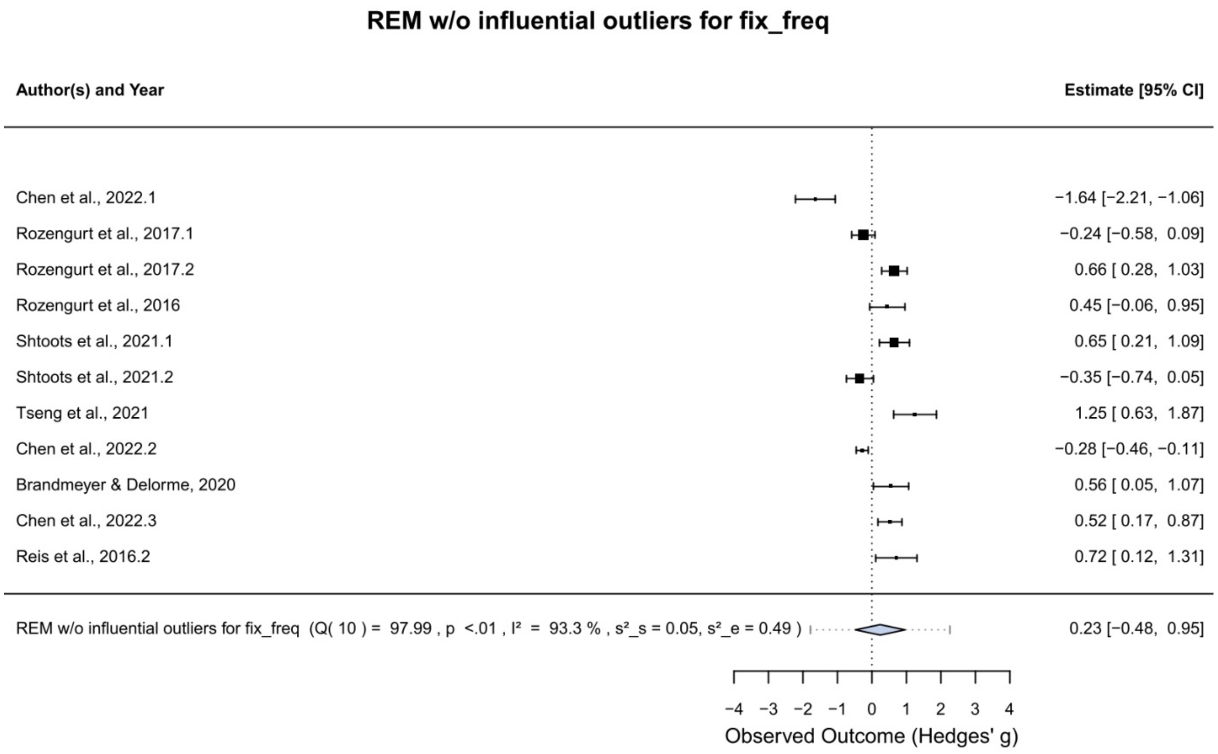


**Fig. S1n** Forest plot for the random-effects model (REM) assessing protocols which used fixed frequencies as feedback without the influential outlier. The black squares denote the mean observed effect sizes, where the size of the square is proportional to the model weight. The confidence interval (95%) is indicated by the whiskers. Diamonds depict the 95%-CI of the REM and their respective prediction intervals (dotted whiskers).

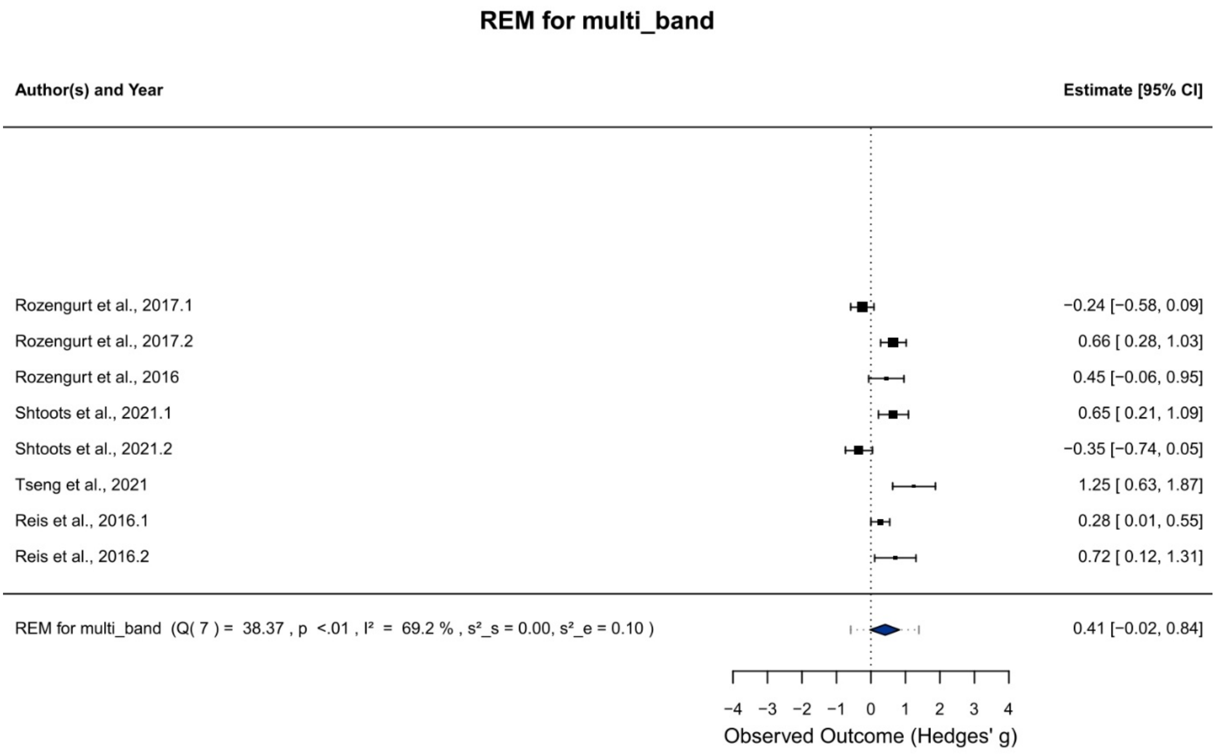


**Fig. S1o** Forest plot for the random-effects model (REM) assessing protocols which used multiple frequency bands as feedback. The black squares denote the mean observed effect sizes, where the size of the square is proportional to the model weight. The confidence interval (95%) is indicated by the whiskers. Diamonds depict the 95%-CI of the REM and their respective prediction intervals (dotted whiskers).

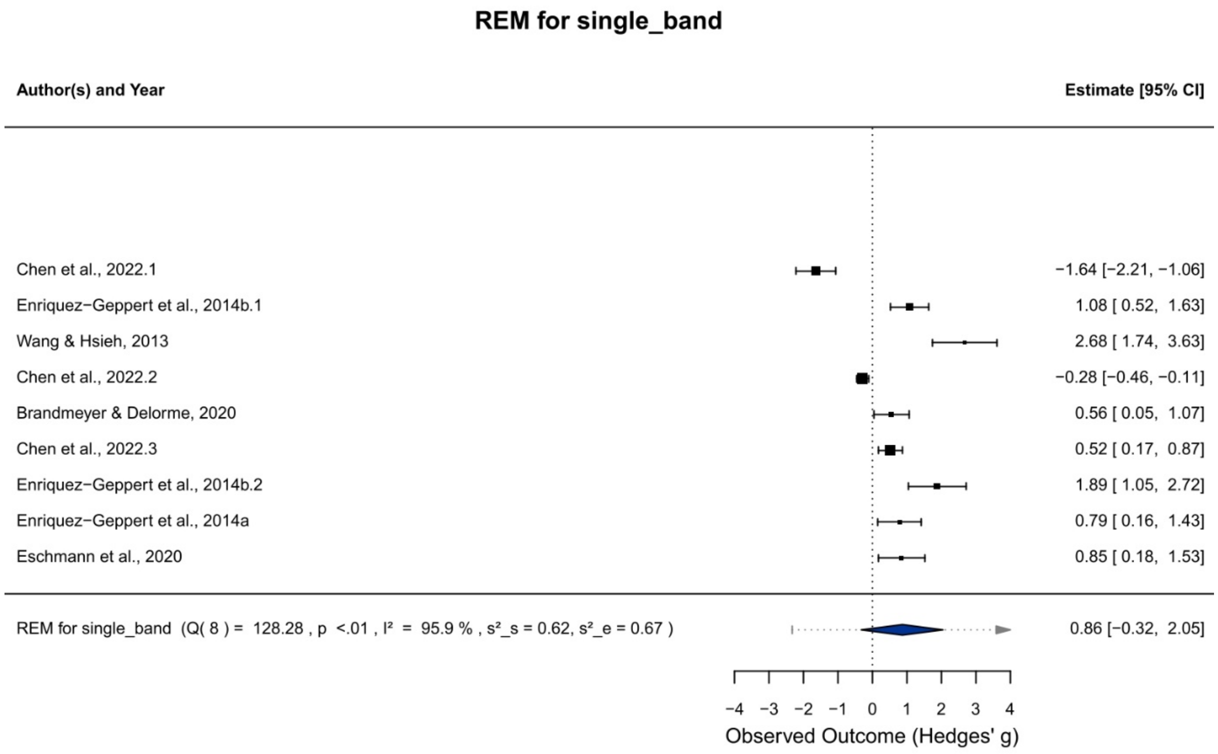


**Fig. S1p** Forest plot for the random-effects model (REM) assessing protocols which used only frontal midline theta as feedback. The black squares denote the mean observed effect sizes, where the size of the square is proportional to the model weight. The confidence interval (95%) is indicated by the whiskers. Diamonds depict the 95%-CI of the REM and their respective prediction intervals (dotted whiskers).

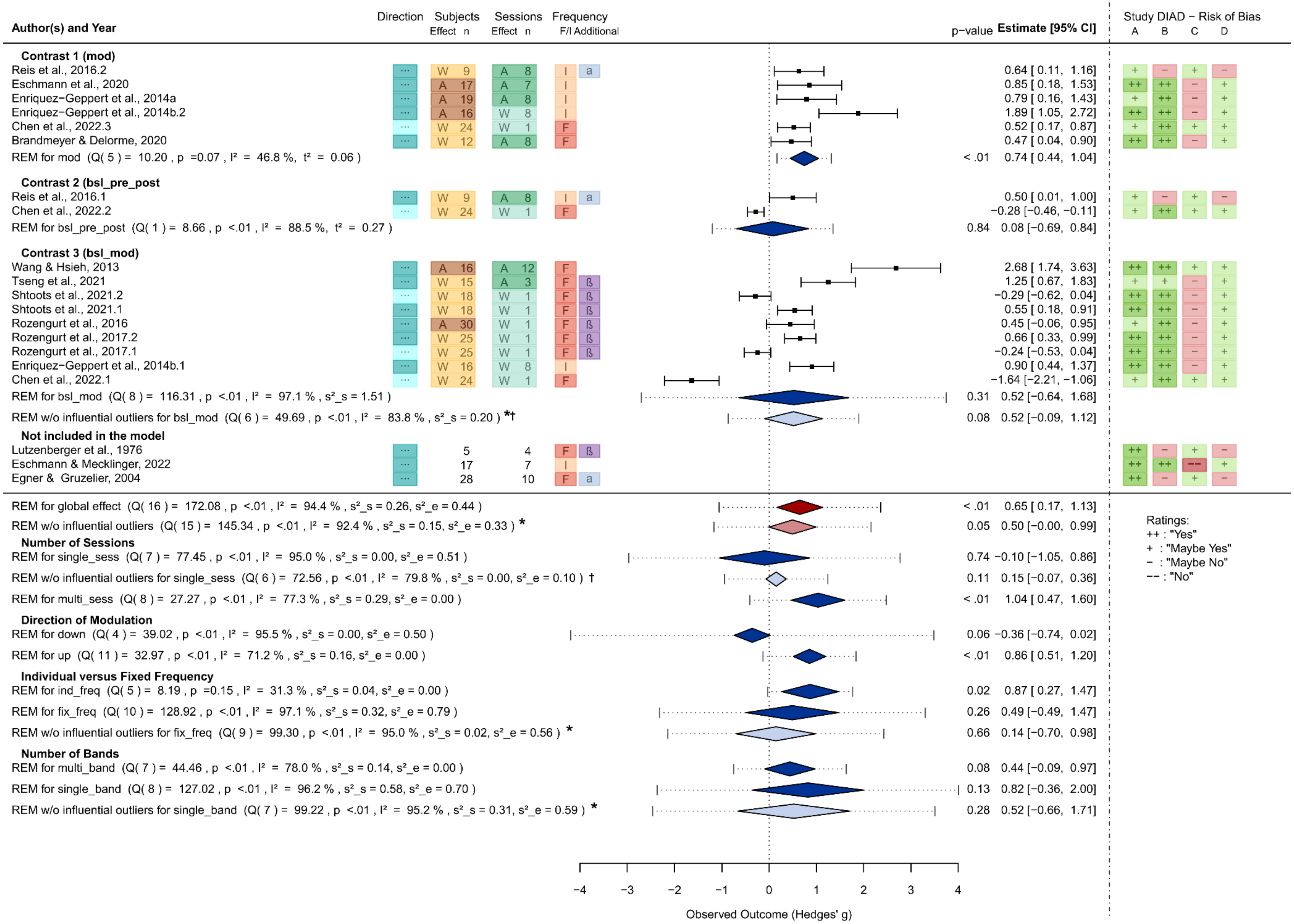
Author(s) and Year
Direction
Subjects Effect n
Sessions Effect n
Frequency F/I Additional
p−value
Estimate [95% CI]
Study DIAD − Risk of Bias
A B C D
Contrast 1 (mod)
Reis et al., 2016.2 0.64 [ 0.11, 1.16]
Eschmann et al., 2020 0.85 [ 0.18, 1.53]
Enriquez−Geppert et al., 2014a 0.79 [ 0.16, 1.43]
Enriquez−Geppert et al., 2014b.2 1.89 [ 1.05, 2.72]
Chen et al., 2022.3 0.52 [ 0.17, 0.87]
Brandmeyer & Delorme, 2020 0.47 [ 0.04, 0.90]
REM for mod (Q( 5 ) = 10.20 , p =0.07 , I² = 46.8 %, t² = 0.06 ) < .01 0.74 [ 0.44, 1.04]
Contrast 2 (bsl_pre_post
Reis et al., 2016.1 0.50 [ 0.01, 1.00]
Chen et al., 2022.2 −0.28 [−0.46, −0.11]
REM for bsl_pre_post (Q( 1 ) = 8.66 , p <.01 , I² = 88.5 %, t² = 0.27 ) 0.84 0.08 [−0.69, 0.84]
Contrast 3 (bsl_mod)
Wang & Hsieh, 2013 2.68 [ 1.74, 3.63]
Tseng et al., 2021 1.25 [ 0.67, 1.83]
Shtoots et al., 2021.2 −0.29 [−0.62, 0.04]
Shtoots et al., 2021.1 0.55 [ 0.18, 0.91]
Rozengurt et al., 2016 0.45 [−0.06, 0.95]
Rozengurt et al., 2017.2 0.66 [ 0.33, 0.99]
Rozengurt et al., 2017.1 −0.24 [−0.53, 0.04]
Enriquez−Geppert et al., 2014b.1 0.90 [ 0.44, 1.37]
Chen et al., 2022.1 −1.64 [−2.21, −1.06]
REM for bsl_mod (Q( 8 ) = 116.31 , p <.01 , I² = 97.1 % , s²_s = 1.51 ) 0.31 0.52 [−0.64, 1.68]
REM w/o influential outliers for bsl_mod (Q( 6 ) = 49.69 , p <.01 , I² = 83.8 % , s²_s = 0.20 ) *† 0.08 0.52 [−0.09, 1.12]
Not included in the model
Lutzenberger et al., 1976 5 4
Eschmann & Mecklinger, 2022 17 7
Egner & Gruzelier, 2004 28 10
REM for global effect (Q( 16 ) = 172.08 , p <.01 , I² = 94.4 % , s²_s = 0.26, s²_e = 0.44 ) < .01 0.65 [ 0.17, 1.13]
REM w/o influential outliers (Q( 15 ) = 145.34 , p <.01 , I² = 92.4 % , s²_s = 0.15, s²_e = 0.33 ) * 0.05 0.50 [−0.00, 0.99]
Number of Sessions
REM for single_sess (Q( 7 ) = 77.45 , p <.01 , I² = 95.0 % , s²_s = 0.00, s²_e = 0.51 ) 0.74 −0.10 [−1.05, 0.86]
REM w/o influential outliers for single_sess (Q( 6 ) = 72.56 , p <.01 , I² = 79.8 % , s²_s = 0.00, s²_e = 0.10 ) † 0.11 0.15 [−0.07, 0.36]
REM for multi_sess (Q( 8 ) = 27.27 , p <.01 , I² = 77.3 % , s²_s = 0.29, s²_e = 0.00 ) < .01 1.04 [ 0.47, 1.60]
Direction of Modulation
REM for down (Q( 4 ) = 39.02 , p <.01 , I² = 95.5 % , s²_s = 0.00, s²_e = 0.50 ) 0.06 −0.36 [−0.74, 0.02]
REM for up (Q( 11 ) = 32.97 , p <.01 , I² = 71.2 % , s²_s = 0.16, s²_e = 0.00 ) < .01 0.86 [ 0.51, 1.20]
Individual versus Fixed Frequency
REM for ind_freq (Q( 5 ) = 8.19 , p =0.15 , I² = 31.3 % , s²_s = 0.04, s²_e = 0.00 ) 0.02 0.87 [ 0.27, 1.47]
REM for fix_freq (Q( 10 ) = 128.92 , p <.01 , I² = 97.1 % , s²_s = 0.32, s²_e = 0.79 ) 0.26 0.49 [−0.49, 1.47]
REM w/o influential outliers for fix_freq (Q( 9 ) = 99.30 , p <.01 , I² = 95.0 % , s²_s = 0.02, s²_e = 0.56 ) * 0.66 0.14 [−0.70, 0.98]
Number of Bands
REM for multi_band (Q( 7 ) = 44.46 , p <.01 , I² = 78.0 % , s²_s = 0.14, s²_e = 0.00 ) 0.08 0.44 [−0.09, 0.97]
REM for single_band (Q( 8 ) = 127.02 , p <.01 , I² = 96.2 % , s²_s = 0.58, s²_e = 0.70 ) 0.13 0.82 [−0.36, 2.00]
REM w/o influential outliers for single_band (Q( 7 ) = 99.22 , p <.01 , I² = 95.2 % , s²_s = 0.31, s²_e = 0.59 ) * 0.28 0.52 [−0.66, 1.71]
Ratings:
++ : "Yes"
+ : "Maybe Yes"
− : "Maybe No"
−− : "No"
−4 −3 −2 −1 0 1 2 3 4
Observed Outcome (Hedges' g)

**Fig. S2a** Forest plot of the multivariate random effect model (REM) including all studies with the contrast-specific analysis depicted below, based on the liberal correction for dependent measurements ($r$ = 0.74; McEvoy et al., 2000): Effects are identified by author, year, and an additional number to distinguish them (whenever several effects were obtained within a single study). Study characteristics with importance to the moderator analyses are displayed in the following way: Direction of modulation: **up (↑) / down (↓)**; number **(n)** of **subjects** in the experimental group and **sessions**; whether the **effect** was measured **within (W)** and **across (A) subjects** or **sessions**; whether a **fixed (F)** or **individual (I) frequency** band was used and if the modulation of an **additional frequency** band was intended. The contrasts of interest (1-3) correspond to the extracted effects: **"mod"** - Start-NF FMT modulation vs. end-NF FMT modulation, **"bsl_pre_post" -** Pre-NF FMT rest vs. post-NF FMT rest and "**bsl_mod**" - Pre-NF FMT rest vs. during-NF FMT modulation. The observed outcomes of FMT modulation are presented as the standardized mean differences (Hedges' g). Means of individual effects are depicted by black squares with solid whiskers which indicate the 95%-Confidence Interval (CI). Diamonds with dotted whiskers depict the 95%-CI of the REMs for the global effects (red), the contrast-specific and moderator effects (blue) and their respective prediction intervals (dotted whiskers). Lighter colours indicate the exclusion of influential outliers. These excluded studies (Chen et al., 2022.1 (**†**) and Wang & Hsieh, 2013(*****)) are identified in the corresponding columns. Study heterogeneity is reported by **$I^2$**. While for univariate models the between-study variance is given by **$\tau^2$**, for multivariate models the between study-variance is indicated by **$\sigma^2$_s** and the within-study variance by **$\sigma^2$_e**. For each reported effect the results of the risk of bias assessment of the corresponding study are presented in the four most right columns: (**A**) Fit between Concept & Operations, (**B**) Clarity of Causal Inference, (**C**) Generalizability of Findings and (**D**) Precision of Outcome Estimation.

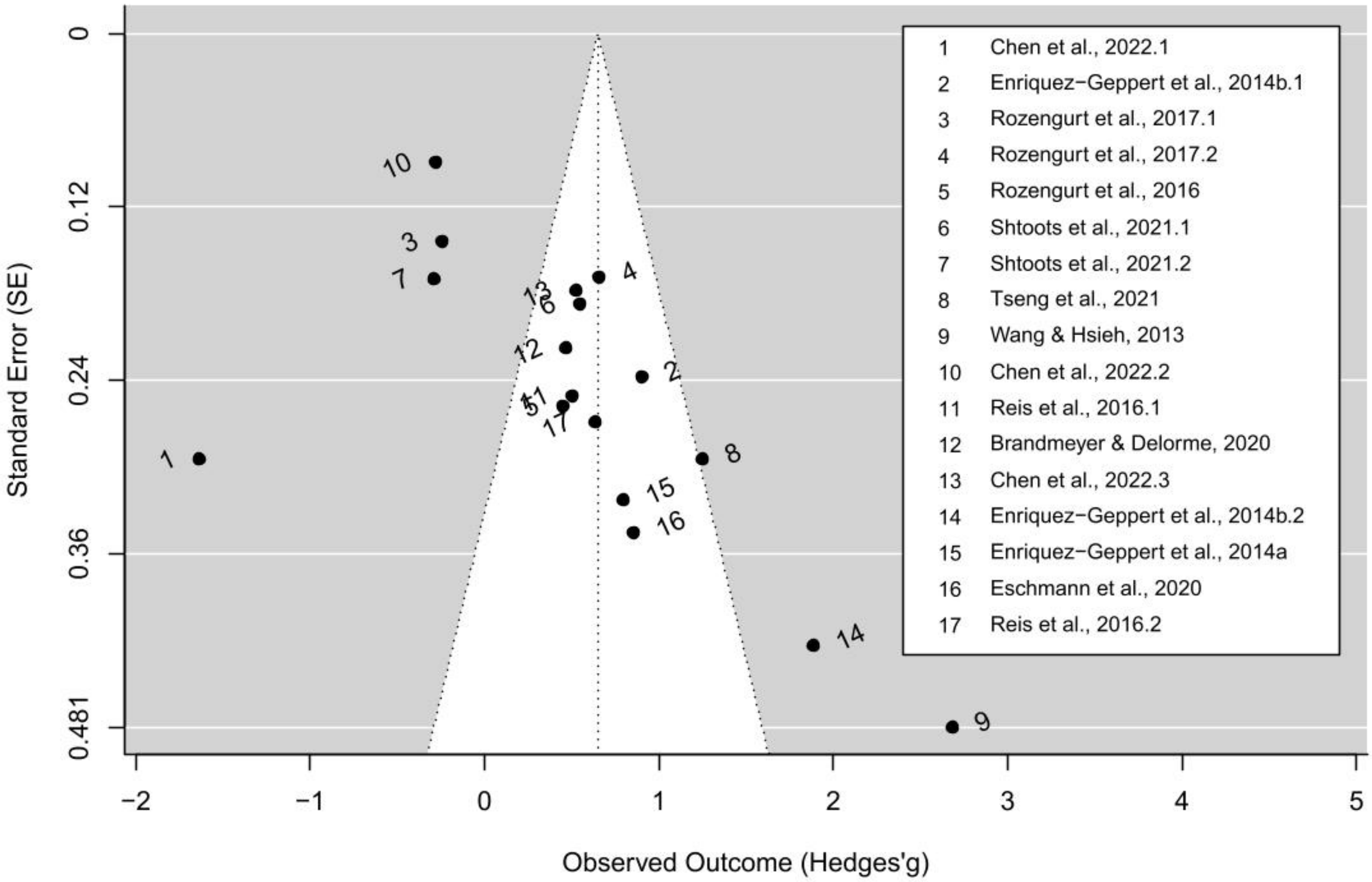


**Fig. S2b** Funnel plot for the assessment of publication bias for the effects included in the quantitative analysis based on the liberal correction for dependent measurements ($r$ = 0.74; McEvoy et al., 2000): Effects are presented by dots. The size of the observed effects (Hedge's $g$) is mapped on the x-axis, whilst the y-axis represents the standard error. The displayed distribution, as well as the significant rank correlation test (Kendall's $\tau$ = 0.37, $p$ = .04) indicate funnel plot asymmetry.

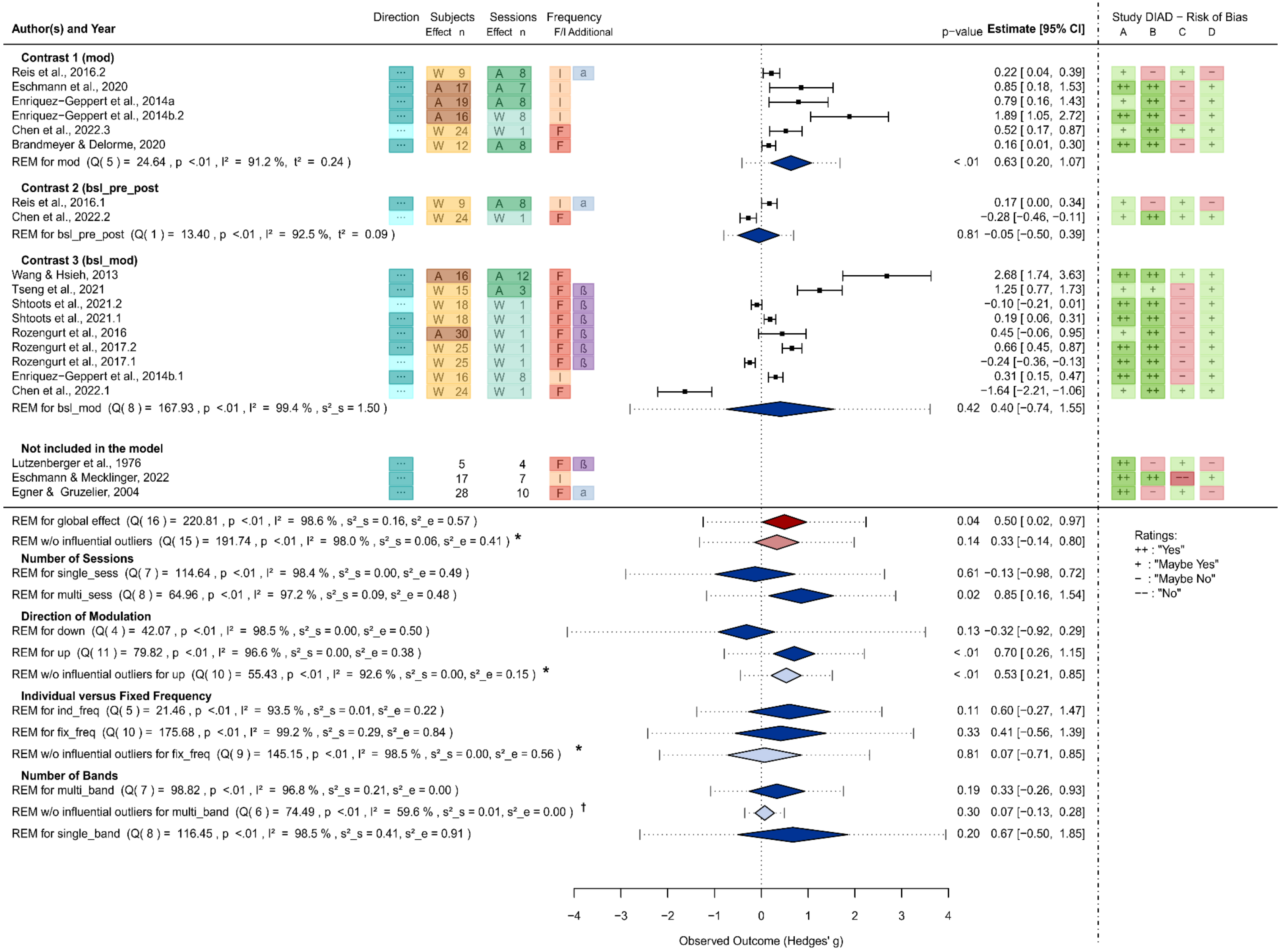
Author(s) and Year
Direction
Subjects Effect n
Sessions Effect n
Frequency F/I Additional
p−value
Estimate [95% CI]
Study DIAD − Risk of Bias
A B C D
Contrast 1 (mod)
Reis et al., 2016.2 — W 9 — A 8 — I a — 0.22 [ 0.04, 0.39] — + − + −
Eschmann et al., 2020 — A 17 — A 7 — I — 0.85 [ 0.18, 1.53] — ++ ++ − +
Enriquez−Geppert et al., 2014a — A 19 — A 8 — I — 0.79 [ 0.16, 1.43] — + ++ − +
Enriquez−Geppert et al., 2014b.2 — A 16 — W 8 — I — 1.89 [ 1.05, 2.72] — ++ ++ − +
Chen et al., 2022.3 — W 24 — W 1 — F — 0.52 [ 0.17, 0.87] — + ++ + +
Brandmeyer & Delorme, 2020 — W 12 — A 8 — F — 0.16 [ 0.01, 0.30] — ++ ++ − +
REM for mod (Q( 5 ) = 24.64 , p <.01 , I² = 91.2 %, τ² = 0.24 ) — < .01 — 0.63 [ 0.20, 1.07]
Contrast 2 (bsl_pre_post)
Reis et al., 2016.1 — W 9 — A 8 — I a — 0.17 [ 0.00, 0.34] — + − + −
Chen et al., 2022.2 — W 24 — W 1 — F — −0.28 [−0.46, −0.11] — + ++ + +
REM for bsl_pre_post (Q( 1 ) = 13.40 , p <.01 , I² = 92.5 %, τ² = 0.09 ) — 0.81 — −0.05 [−0.50, 0.39]
Contrast 3 (bsl_mod)
Wang & Hsieh, 2013 — A 16 — A 12 — F — 2.68 [ 1.74, 3.63] — ++ ++ + +
Tseng et al., 2021 — W 15 — A 3 — F ß — 1.25 [ 0.77, 1.73] — + + − +
Shtoots et al., 2021.2 — W 18 — W 1 — F ß — −0.10 [−0.21, 0.01] — ++ ++ − +
Shtoots et al., 2021.1 — W 18 — W 1 — F ß — 0.19 [ 0.06, 0.31] — ++ ++ − +
Rozengurt et al., 2016 — A 30 — W 1 — F ß — 0.45 [−0.06, 0.95] — + ++ − +
Rozengurt et al., 2017.2 — W 25 — W 1 — F ß — 0.66 [ 0.45, 0.87] — ++ ++ − +
Rozengurt et al., 2017.1 — W 25 — W 1 — F ß — −0.24 [−0.36, −0.13] — ++ ++ − +
Enriquez−Geppert et al., 2014b.1 — W 16 — W 8 — I — 0.31 [ 0.15, 0.47] — ++ ++ − +
Chen et al., 2022.1 — W 24 — W 1 — F — −1.64 [−2.21, −1.06] — + ++ + +
REM for bsl_mod (Q( 8 ) = 167.93 , p <.01 , I² = 99.4 % , s²_s = 1.50 ) — 0.42 — 0.40 [−0.74, 1.55]
Not included in the model
Lutzenberger et al., 1976 — 5 — 4 — F ß — ++ − + −
Eschmann & Mecklinger, 2022 — 17 — 7 — I — ++ ++ −− +
Egner & Gruzelier, 2004 — 28 — 10 — F a — ++ − + −
REM for global effect (Q( 16 ) = 220.81 , p <.01 , I² = 98.6 % , s²_s = 0.16, s²_e = 0.57 ) — 0.04 — 0.50 [ 0.02, 0.97]
REM w/o influential outliers (Q( 15 ) = 191.74 , p <.01 , I² = 98.0 % , s²_s = 0.06, s²_e = 0.41 ) * — 0.14 — 0.33 [−0.14, 0.80]
Number of Sessions
REM for single_sess (Q( 7 ) = 114.64 , p <.01 , I² = 98.4 % , s²_s = 0.00, s²_e = 0.49 ) — 0.61 — −0.13 [−0.98, 0.72]
REM for multi_sess (Q( 8 ) = 64.96 , p <.01 , I² = 97.2 % , s²_s = 0.09, s²_e = 0.48 ) — 0.02 — 0.85 [ 0.16, 1.54]
Direction of Modulation
REM for down (Q( 4 ) = 42.07 , p <.01 , I² = 98.5 % , s²_s = 0.00, s²_e = 0.50 ) — 0.13 — −0.32 [−0.92, 0.29]
REM for up (Q( 11 ) = 79.82 , p <.01 , I² = 96.6 % , s²_s = 0.00, s²_e = 0.38 ) — < .01 — 0.70 [ 0.26, 1.15]
REM w/o influential outliers for up (Q( 10 ) = 55.43 , p <.01 , I² = 92.6 % , s²_s = 0.00, s²_e = 0.15 ) * — < .01 — 0.53 [ 0.21, 0.85]
Individual versus Fixed Frequency
REM for ind_freq (Q( 5 ) = 21.46 , p <.01 , I² = 93.5 % , s²_s = 0.01, s²_e = 0.22 ) — 0.11 — 0.60 [−0.27, 1.47]
REM for fix_freq (Q( 10 ) = 175.68 , p <.01 , I² = 99.2 % , s²_s = 0.29, s²_e = 0.84 ) — 0.33 — 0.41 [−0.56, 1.39]
REM w/o influential outliers for fix_freq (Q( 9 ) = 145.15 , p <.01 , I² = 98.5 % , s²_s = 0.00, s²_e = 0.56 ) * — 0.81 — 0.07 [−0.71, 0.85]
Number of Bands
REM for multi_band (Q( 7 ) = 98.82 , p <.01 , I² = 96.8 % , s²_s = 0.21, s²_e = 0.00 ) — 0.19 — 0.33 [−0.26, 0.93]
REM w/o influential outliers for multi_band (Q( 6 ) = 74.49 , p <.01 , I² = 59.6 % , s²_s = 0.01, s²_e = 0.00 ) † — 0.30 — 0.07 [−0.13, 0.28]
REM for single_band (Q( 8 ) = 116.45 , p <.01 , I² = 98.5 % , s²_s = 0.41, s²_e = 0.91 ) — 0.20 — 0.67 [−0.50, 1.85]
Ratings:
++ : "Yes"
+ : "Maybe Yes"
− : "Maybe No"
−− : "No"
−4 −3 −2 −1 0 1 2 3 4
Observed Outcome (Hedges' g)

**Fig. S3a** Forest plot of the multivariate random effect model (REM) including all studies with the contrast-specific analysis depicted below, based on the conservative correction for dependent measurements (*r* = 0.97; McEvoy et al., 2000): Effects are identified by author, year, and an additional number to distinguish them (whenever several effects were obtained within a single study). Study characteristics with importance to the moderator analyses are displayed in the following way: Direction of modulation: **up (↑) / down (↓)**; number **(n)** of **subjects** in the experimental group and **sessions**; whether the **effect** was measured **within (W)** and **across (A) subjects** or **sessions**; whether a **fixed (F)** or **individual (I) frequency** band was used and if the modulation of an **additional frequency** band was intended. The contrasts of interest (1-3) correspond to the extracted effects: **"mod"** - Start-NF FMT modulation vs. end-NF FMT modulation, **"bsl_pre_post" -** Pre-NF FMT rest vs. post-NF FMT rest and "**bsl_mod**" - Pre-NF FMT rest vs. during-NF FMT modulation. The observed outcomes of FMT modulation are presented as the standardized mean differences (Hedges' g). Means of individual effects are depicted by black squares with solid whiskers which indicate the 95%-Confidence Interval (CI). Diamonds with dotted whiskers depict the 95%-CI of the REMs for the global effects (red), the contrast-specific and moderator effects (blue) and their respective prediction intervals (dotted whiskers). Lighter colours indicate the exclusion of influential outliers. These excluded studies (Chen et al., 2022.1 (**†**) and Wang & Hsieh, 2013(*****)) are identified in the corresponding columns. Study heterogeneity is reported by $\mathbf{I^2}$. While for univariate models the between-study variance is given by $\boldsymbol{\tau^2}$, for multivariate models the between study-variance is indicated by $\boldsymbol{\sigma^2\_s}$ and the within-study variance by $\boldsymbol{\sigma^2\_e}$. For each reported effect the results of the risk of bias assessment of the corresponding study are presented in the four most right columns: (**A**) Fit between Concept & Operations, (**B**) Clarity of Causal Inference, (**C**) Generalizability of Findings and (**D**) Precision of Outcome Estimation.

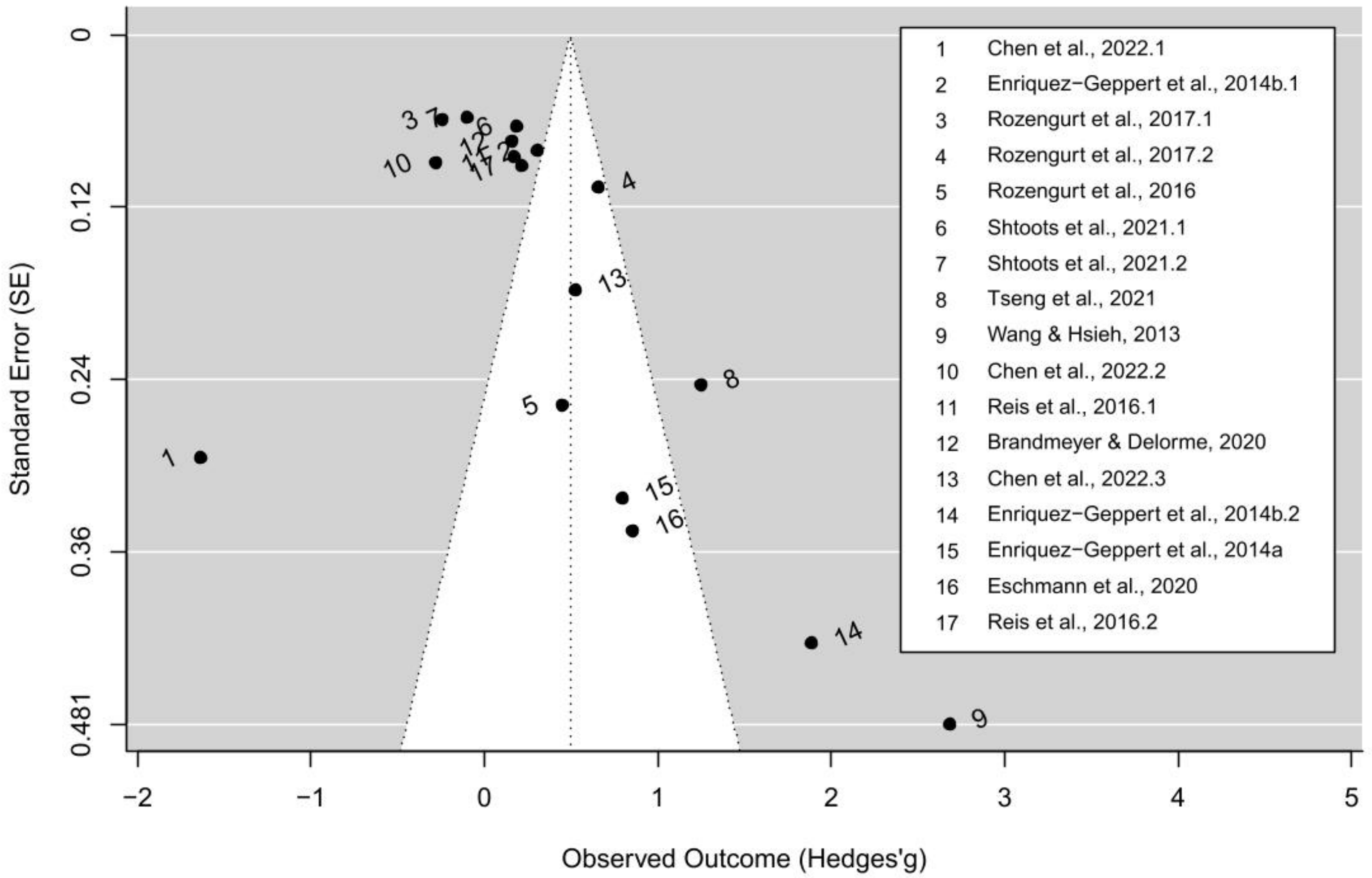


**Fig. S3b** Funnel plot for the assessment of publication bias for the effects included in the quantitative analysis, based on the conservative correction for dependent measurements ($r$ = 0.97; McEvoy et al., 2000): Effects are presented by dots. The size of the observed effects (Hedge's $g$) is mapped on the x-axis, whilst the y-axis represents the standard error. The displayed distribution as well as the significant rank correlation test (Kendall's $\tau$ = 0.38, $p$ = .03) indicate funnel plot asymmetry
